# Long-Term Ecological Baselines and Critical Thresholds in Ombrotrophic Peatlands of Europe: Implications for Restoration Strategies

**DOI:** 10.1101/2025.05.16.654541

**Authors:** Mateusz Draga, Mariusz Gałka, Klaus-Holger Knorr, Stephan Glatzel, Bogdan H. Chojnicki, Christian Fritz, Vincent E.J. Jassey, Radosław Juszczak, Hanna Meyer, Bjorn J.M. Robroek, Carrie L. Thomas, Mariusz Lamentowicz

## Abstract

Maintaining appropriate peatland hydration, particularly through the regulation of the depth to the water table (DWT), is crucial for peatland conservation, restoration, and the mitigation of greenhouse gas emissions. In this study, we assess the long-term ecological impact of hydrological changes, primarily induced by drainage, on ombrotrophic peatlands across Europe. Our analysis is based on novel palaeoecological data from seven peat cores collected from sites that have experienced varying degrees of anthropogenic disturbance. We reconstructed historical DWT fluctuations using plant macrofossil and testate amoeba analyses at high resolution. By applying Threshold Indicator Taxa Analysis (TITAN), we identified species-specific and community-level response thresholds to changes in water level. This approach revealed two distinct change points: the first, at approximately 7 cm DWT, corresponds to hydrological conditions favourable for moisture-dependent *Sphagnum* species, while the second, near 22 cm DWT, is associated with more drought-adapted taxa and signals ecosystem degradation. The interval between these points represents a transition zone between optimal and suboptimal conditions for peatland functioning. An additional TITAN analysis aimed at identifying the timing of the most significant ecological changes indicates that peatland degradation intensified over the past two centuries and has accelerated in recent decades. Our findings also show that even after hydrological restoration, microbial and plant communities often remain distinct from those in undisturbed peatlands. This underscores the importance of preserving sites that still retain near-natural conditions. Based on our results (and consistent with previous studies) we recommend maintaining a DWT of approximately 10 cm below the surface as an optimal target for both peatland conservation and restoration. Such conditions not only support ecological integrity but are also associated with reduced greenhouse gas emissions and higher peat accumulation rates, reinforcing the role of ombrotrophic peatlands as long-term carbon sinks.

## 1. Introduction

With ongoing climate change, peatlands remain one of our most valuable natural allies in efforts to mitigate or offset their impacts (Strack et al. 2022). Although they occupy only a small fraction of Earth’s surface, peatlands support a wide range of essential ecosystem services and serve as important regulators of environmental processes (Kimmel and Mander 2010, Loisel et al. 2021a). Intact peatlands contribute to groundwater stabilisation, enhance resilience to drought (Lennartz and Liu 2019, Taufik et al. 2020), exert localised cooling effects (Worrall et al. 2020), and help mitigate the risk of wildfires (Kettridge et al. 2017, Taufik et al. 2022). Moreover, peatlands often form biodiversity-rich microhabitats within otherwise intensively managed forest or agricultural landscapes. In addition to these ecological functions, peatlands play a pivotal role in the global carbon cycle. They are estimated to store between 30% and 44% of the world’s soil carbon (Parish et al. 2008, Joosten et al. 2016), an amount comparable to the carbon currently present in the atmosphere (Yu 2012, Dargie et al. 2017). These vast carbon reservoirs have developed over tens of thousands of years through the slow accumulation of plant biomass. To slow the pace of global warming, it is imperative that this organic carbon remains stored in the peat (Strack et al. 2022). In this context, it is particularly alarming that many peatlands are now degraded, characterised by lowered water tables and a notable absence of active peat-forming processes (Page and Baird 2016, Swindles et al. 2019, Tanneberger et al. 2021a), due to decades – even centuries–long pressure from human activity on these ecosystems (Loisel et al. 2021b). As a result of these changes, such sites not only lose their ability to absorb carbon from the atmosphere but, due to ongoing peat oxygenation, begin to emit significant amounts of greenhouse gases back into the atmosphere (Leifeld et al. 2019, Doelman et al. 2023). Thus, considering both the benefits of well-preserved peatlands and the dangers posed by degraded ones, it is not surprising that increasing efforts are dedicated annually to various peatland protection and restoration projects (Apori et al. 2022). Regrettably, despite the existence of numerous restoration methods, such as revegetation, drain blocking, rewetting, ditch blocking, tree harvesting, herbicide spraying, and topsoil removal, peatland restoration remains a complex challenge (Artz et al. 2018, Zak and McInnes 2022, Stachowicz et al. 2023).

The functioning of peatlands largely depends on well-balanced and relatively stable hydrological conditions. Determining the depth to the water table (DWT) (Baird and Low 2022) that supports both the functioning of intact peatlands and the regeneration of degraded ones is, however, a complex task (Lennartz and Liu 2019, Austin et al. 2025). That said, it should be noted that the DWT is not a fixed value and often varies significantly across seasons (Breeuwer et al. 2009). Despite these natural fluctuations, many peatlands have, in recent decades, experienced a sustained lowering of the water table, leading to more frequent drought conditions. This has, in turn, negatively affected the ecology of numerous species that are highly specialised and confined to peatland habitats. Over time, hydrophilic plant species typical of intact ecosystems may be replaced by species better adapted to lower water table depths and drier conditions (Jassey et al. 2018), initiating a successional shift toward vascular plant dominance that ultimately accelerates peatland degradation. Thus, ongoing plant species turnover on peatlands is usually a sign of rapidly progressing ecosystem degradation (Dieleman et al. 2015). Moreover, such turnover events do not solely affect plant communities, as such disturbances will also result in severe changes in invertebrate and microorganism species composition (Lamentowicz and Mitchell 2005, Swindles et al. 2016). However, because substantial shifts in community composition are typically well preserved in peat layers, these records offer researchers a valuable opportunity to reconstruct historical changes in water table depth (DWT) and assess their long-term effects on species composition (Lamentowicz and Mitchell 2005, Chambers et al. 2012), thereby providing a stronger basis for informing ecosystem restoration efforts.

Linking changes in plant species composition from macrofossil data to proxies on concomitant changes in water table depth may provide insight into the historical response of the plant community to drought. (Booth et al. 2006, Swindles et al. 2010). The results can then be analysed to determine the critical DWT threshold beyond which peatland degradation begins (Bragazza 2008). Identifying such thresholds is crucial for effective peatland restoration and conservation efforts. While, in most cases, predicting the value of a change point is a complicated task (Scheffer et al. 2009) whose estimation may be tried through field studies (Jassey et al. 2018), fossil data provides us with a unique opportunity to estimate this parameter directly from the degradation events that happened in the past. Additionally, since paleodata can contain information spanning hundreds or even thousands of years, such data can provide researchers with hundreds of unique records on the relationship between plant community and peatland hydration to analyse. Moreover, these records are chronologically ordered, offering valuable insights into temporal changes and succession. Such an approach was utilised in the work of Lamentowicz et al. (2019), who estimated a plant community tipping point for the DWT value analysing the fossil records collected from seven *Sphagnum*-dominated peatlands (raised bogs and poor fens) in Poland, with samples representing the palaeoecological data from the past 2000 years.

The primary data source for our analysis is fossil records from seven ombrotrophic peatlands of diverse characteristics, spanning five European countries. Each site provides an average history spanning the past 2000 years. Notably, each site possesses a unique history, characterised by varying ages, preservation states, degradation levels, and exploitation histories. All of these sites have recently participated in restoration programs. This study aims to determine the critical transition zone for hydrology in European ombrotrophic peatlands and identify the time during which the most significant ecological change associated with drainage and peat harvesting was observed. According to the former palaeoecological and GHG observational studies, we tested the hypothesis that the critical water table threshold is located at a depth of ca. 10 cm and long-term shifts from this value significantly affect vegetation. Unlike former research, we focused on the disturbance time window and the context of the change point and ecological baselines related to the recent peatland land-use change.

## 2 Methods

### 2.1. Study sites

The research is based on the data obtained from the analysis of seven different peat profiles, gathered from primarily various *Sphagnum-dominated* peatlands located in Europe: Amtsvenn-Hündfelder Moor (Germany), Bagno Kusowo (Poland), Drebbersches Moor (Germany), Fochteloerveen (Netherland), Pichlmeier Moor (Austria), Pürgschachen Moor (Austria) and Store Mosse (Sweden) (Fig.1). The selected sites encompass peatlands with diverse histories and varying degrees of preservation.

**Figure. 1.**
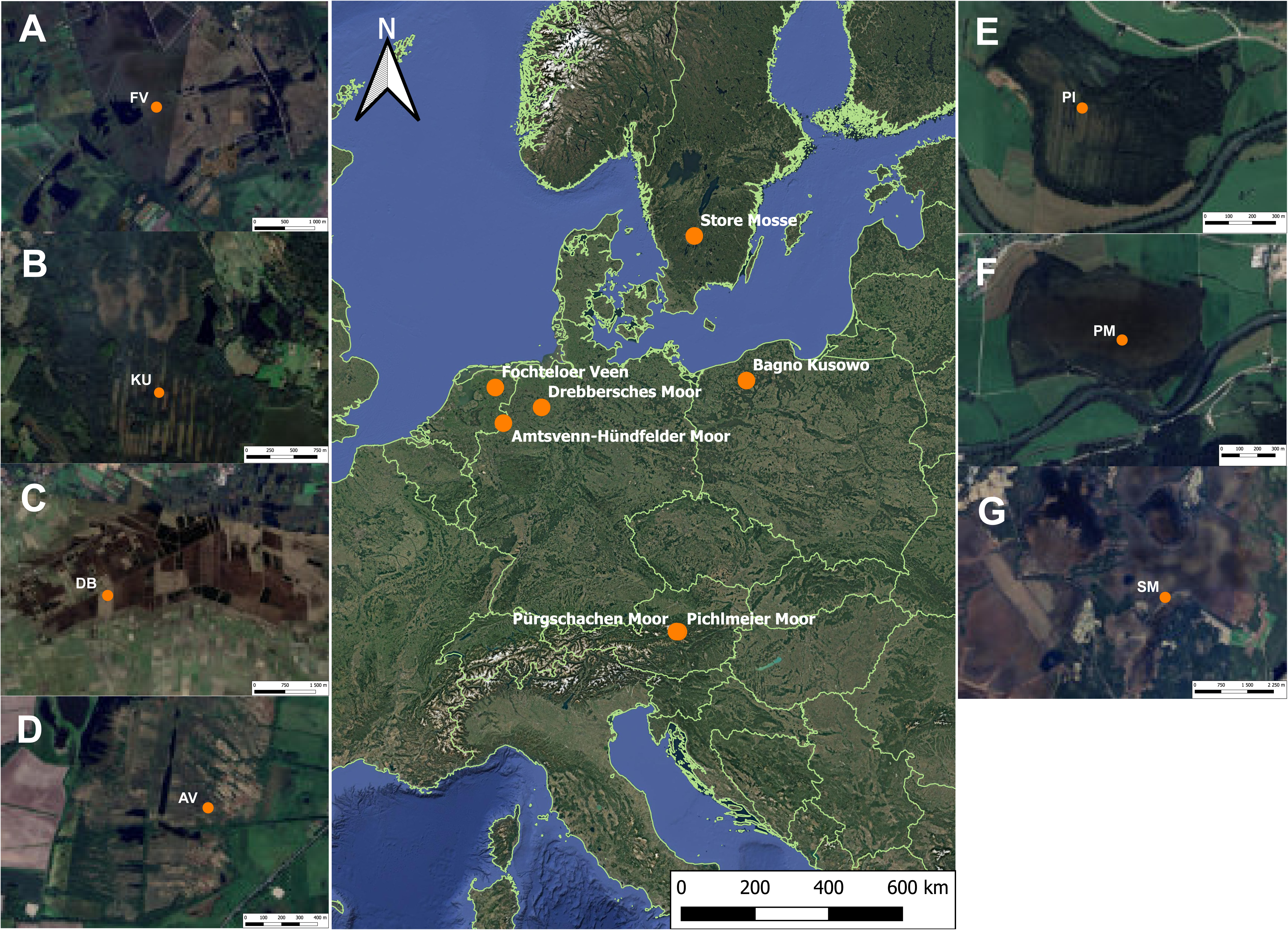
Locations of the seven sites from which peat cores were collected and subsequently analysed. FV - Fochteloërveen (Netherland), KU - Bagno Kusowo (Poland), DB - Drebbersches Moor (Germany), AV - Amtsvenn-Hündfelder Moor (Germany), PI - Pichlmeier Moor (Austria), PM - Pürgschachen Moor (Austria), SM - Store Mosse (Sweden).

**Figure. 2.**
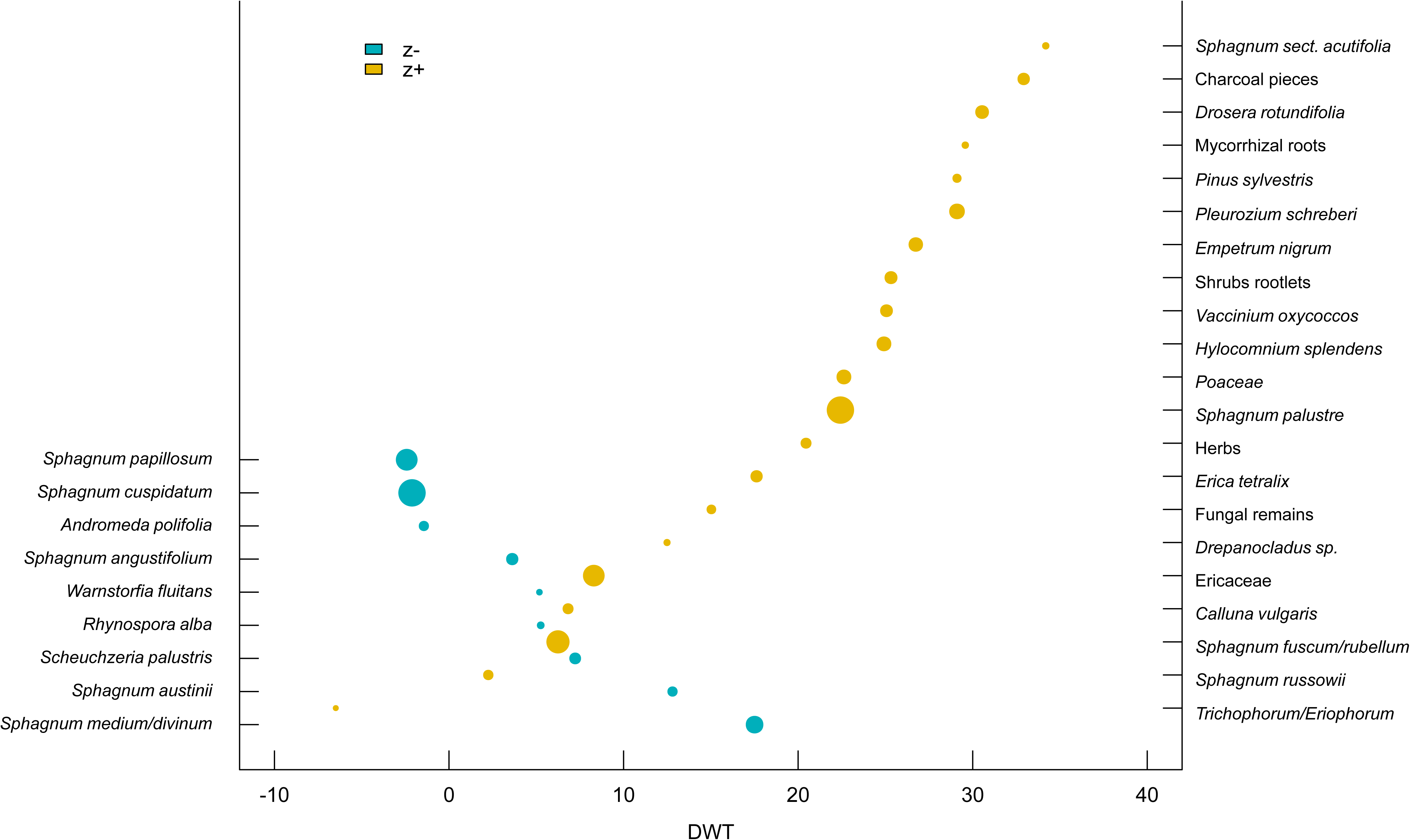
Plot showing taxon-specific change points along the DWT gradient for taxa associated with wet (z-) and dry (z+) conditions. Positive values of the DWT parameter indicate a water table level below the surface, while negative values represent water accumulation above ground level.

#### Amtsvenn-Hündfelder Moor (Germany)

The Amtsvenn-Hündfelder Moor, located in North Rhine-Westphalia, Germany, near the Dutch border, has a long history of human impact. Significant peat extraction and agricultural reclamation began in the early 19th century, primarily for buckwheat cultivation. Despite these activities, the area was once renowned for its diverse lagg vegetation. However, in the 20th century, industrial peat extraction and large-scale reclamation efforts caused profound changes in the landscape. The construction of drainage ditches and peat cutting led to severe peat degradation and surface drying, resulting in vegetation shifts. Today, the site is protected as part of the Natura 2000 network.

#### Bagno Kusowo (Poland)

Bagno Kusowo is a raised bog located in the northwestern part of Poland and is recognised as one of the best examples of a Baltic-type raised bog in this country. Extensive peat extraction began in the late 19th century and continued into the early 20th century. By the early 1960s, the bog had been completely drained to facilitate further peat extraction, with most activity concentrated in the southern part of the peatland, which is now undergoing restoration. The northern section has remained waterlogged despite the drainage, likely preserving much of its original ecological characteristics. The site is protected as part of a nature reserve and the Natura 2000 network.

#### Drebbersches Moor (Germany)

Drebbersches Moor is a raised bog located in Lower Saxony in Germany that is a part of the larger raised bog area of Großes Moor. Before its allocation as a protected nature area, the site was used for buckwheat cultivation since the late 18th century and consequently subjected to surficial drainage and controlled burning. Dams have been built to reduce water losses to adjacent peat harvesting areas surrounding the site. Yet, the site presumably still loses water to surrounding drainage structures within the peat harvesting area. In summer 2003, there was a fire at the site, which mainly affected the surface peat. The site is characterised by shallow water table levels, reaching near-surface levels during the wettest times, yet still too dry for raised bog regeneration.

#### Fochteloërveen (Netherlands)

Fochteloërveen is a cutover bog located in the northern Netherlands, characterised by an accumulated peat layer reaching depths of up to 3 meters. Historically, the site has been impacted by various disturbances, including fires, droughts, occasional flooding, and agricultural activities. Since 2000, it has been undergoing an intensive restoration process focused on large-scale rewetting efforts, which have resulted in partial moss regeneration. Despite these measures, Fochteloer Veen’s future remains threatened due to continued nutrient accumulation and recurring drought events. The entire area is designated as a national reserve and is protected under the Natura 2000 program.

#### Pichlmeier Moor (Austria)

Pichlmaier Moor is a bog located in the Austrian Enns valley in the eastern Alps, approximately 5 km east of Pürgschachen Bog. In the past, the area was extensively drained for peat extraction, as evidenced by three heavily degraded vegetation ridges, between which peat was removed. The excavated sections between these ridges have been rewetted since the early 1990s, with some areas of open water in the centre of the ditches. Currently, the site is undergoing natural vegetation succession, forming floating *Sphagnum*-dominated peat mats in the ditches. Core samples for this research were collected from the edge of the ridge.

#### Pürgschachen Moor (Austria)

Pürgschachen Moor is part of a bog complex in central Austria, situated in the Enns Valley of the eastern Alps, about 5 km west of Pichlmaier Moor. It is the largest relatively undisturbed raised valley bog in the Alps, covering an area of 62 hectares. It is characterised by an intact treeless central area dominated by native peatland vegetation. However, the bog shows signs of degradation as the advancing succession of *Betula pubescens* and *Pinus mugo* is observed.

#### Store Mosse (Sweden)

The Store Mosse National Park is the largest pristine peatland expanse in southern Sweden. The areas adjacent to this peatland were historically subjected to intensive peat extraction. The whole area is currently undergoing restoration, along with several other peatlands in Sweden, as part of the Addmire Project initiated in 2015. Store Mosse is characterised by well-preserved *Sphagnum* vegetation with a lawn-hollow structure. Declared a national park in 1982, the area is legally protected and included in the Natura 2000 programme.

### 2.2 Coring, subsampling and depth to the water table reconstruction

The cores from each site were sampled in 2022-2023 using a Wardenaar sampler (Wardenaar 1987) and an Instorf corer to recover a monolith from the top 1 meter. The samples were transported to the laboratory at the Adam Mickiewicz University, Poznań, Poland, and stored in the cold room (constant temperature +4C). Each core was subsampled for individual proxies, dated, and analysed at various resolutions. Pollen, microscopic charcoal, and testate amoebae were sampled at 5-cm intervals, whereas plant macrofossils and macroscopic charcoal were analysed continuously at 1-cm intervals.

The absolute chronology for each core was based upon 39 ^14^C AMS dates provided by the Poznań Radiocarbon Laboratory (Poland) (Supplementary Table 1). The age-depth model was calculated using the OxCal 4.3 software (Bronk Ramsey 1995) applying the P_Sequence function with parameters: k0=0.6, log10(k/k0)=1, and interpolation = 1 cm (Bronk Ramsey 2008, Ramsey and Lee 2013). The IntCal13 atmospheric curve was used as the calibration dataset (Reimer et al. 2013).

High-resolution (1-cm peat slices of approximately 10 cm^3^ volume, in contiguous samples) plant macrofossil analysis was used to reconstruct local plant succession. The samples were washed and sieved under a warm water current over 0.20 mm mesh screens. The total of vegetative plant remains remaining on the sieve after rinsing the peat was set to 100%. The percentage of individual vascular plants and mosses was then estimated to the nearest 5%. The fossil carpological remains and vegetative fragments (leaves, rootlets, epidermis) were identified using identification keys (Smith 2004, Mauquoy and van Geel 2007) and compared to recently collected specimens. See Gałka et al. (2019) for more details on the methods used for plant macrofossil analysis of peat.

Testate amoeba (TA) material (2 cm^3^ in volume) was sampled from the same depths as the microscopic charcoal analyses. Peat samples were washed under 0.3-mm sieves following the method described by Booth et al. (2010). TAs were analysed under a light microscope between 200× and 400× magnification, with a minimum of 100 tests per sample whenever possible (Payne and Mitchell 2008). Several keys and taxonomic monographs (Grospietsch 1958, Ogden and Hedley 1980, Meisterfeld 2001a, 2001b, Clarke 2003, Mazei and Tsyganov 2006) as well as internet resources (Siemensma 2019) were used to achieve the highest possible taxonomic resolution. Quantitative reconstruction of the TA-based DWT was performed in C2 software (Juggins 2003), using a transfer function approach applying the tolerance-downweighted weighted averaging model (Juggins and Birks 2012) and it was based on the European training set (Amesbury et al. 2016). Diagrams with palaeoecological proxy data were plotted using C2 software (Juggins 2003). DataGraph was used (MacAskill 2012) to draw the synthesis figure.

### 2.3 TITAN threshold analysis

Two threshold indicator taxa analysis (TITAN) were conducted following the method described by Baker and King (2010) to identify DWT and temporal ranges associated with species turnover and species-specific change points in response to DWT fluctuations. TITAN is a statistical method that estimates both taxon-specific and community-level threshold values from ecological data, even for the data characterised by low occurrence for the number of different taxa - a common challenge in ecological studies (King and Baker 2014). During the analysis, each taxon is classified into one of two groups: taxa negatively responding (z-), and taxa positively responding (z+) to increases in the analysed parameter. The community threshold is calculated separately for the z- and z+ group based on the highest Sum(z) peak values along the parameter gradient. The accuracy of the results for each taxon is evaluated with purity and reliability scores. Purity indicates the proportion of times a taxon was assigned to the final group, while reliability represents the proportion of times a taxon achieved a p-value < 0.05 across bootstrap replicates. Although charcoal pieces are not taxa, they were also included in the analysis, as their presence may provide valuable insights into the potential relationship between extremely low DWT values and fire occurrences.

The first TITAN analysis assessed the influence of historical DWT levels on the presence of taxa recorded in the macrofossil data. For this purpose, data from all study sites were pooled and analysed collectively. Thresholds were calculated for all taxa that appeared at least three times in the entire dataset. The second TITAN analysis investigated the impact of time (years) on the presence of different taxa to identify potential community-level species turnover over the last few centuries. Data from six sites were used for this analysis, as the oldest layers from one site (Store Mosse) dated only to the mid-20th century. Additionally, since the analysis focused on the past few centuries, only samples from 1500 to the present were included. Furthermore, as in the previous analysis, only taxa occurring at least three times were included.

For all the calculations the R program (R Core Team 2023) was used, for which the R Studio environment (Posit team 2023) was utilized. During the data preparation and cleaning steps of the study, the dplyr package (Hadley et al. 2023) was used, while the threshold indicator taxa analysis was performed using the Titan2 package (Baker et al. 2015).

## 3. Results

### 3.1. Age-depth models and major change in peat-forming plants

#### Age-depth models and major change in peat-forming plants

Radiocarbon dating indicated that the peat profiles varied considerably in age, ranging from just a few decades at Store Mosse to over 5,000 years at Pichlmeier Moor, with the majority dating back at least 1,500 years (Table 1, Supplementary Table 1). The resulting datasets, which documented historical changes in DWT and plant species occurrences, were combined and used in subsequent analyses focusing on the recent disturbances related to drainage and peat harvesting. The full set of the age-depth models is presented in the Supplementary Material (Supplementary Figure 1-7), complete plant macrofossils diagrams are presented in the Supplementary Material (Supplementary Figure 8-14), complete testate amoebae diagrams are presented in the Supplementary Material (Supplementary Figure 15-21).

**Table 1.** Overview of all sites from which peat cores were extracted, including the country of origin, precise location of the peat core extraction site, brief usage history, state of preservation, age range of analysed material, and the number of analysed layers.

| Site name | Country | Location | History of use | State of preservation | Time range of a sample | No. of analysed samples |
| --- | --- | --- | --- | --- | --- | --- |
| Amtsvenn-Hündfelder Moor | Germany | 52°10'31.3''N,<br>6°57'19.5''E | Drainage,<br>mining | Poor/Rewetting | -307 - 1982 | 89 |
| Bagno Kusowo | Poland | 53°48'45''N,<br>16°35'11.8''E | Drainage,<br>mining | Moderate/Rewetting | 289 - 2020 | 99 |
| Drebbersches Moor | Germany | 52°41'59.8''N,<br>8°21'34.5''E | Drainage,<br>mining | Poor/Rewetting | 637 - 2018 | 99 |
| Fochteloer Veen | Netherlands | 52°59'44.1''N,<br>6°23'39.3''E | Drainage,<br>mining | Poor/Rewetting | 806 - 2020 | 92 |
| Pichlmeier Moor | Austria | 47°34'49.3''N,<br>14°24'55.3''E | Drainage,<br>mining | Moderate/rewetting | -3370 - 2016 | 45 |
| Pürgschachen Moor | Austria | 47°34'50.0''N,<br>14°20'50.0''E | Drainage | Moderate | -339 - 2018 | 200 |
| Store Mosse | Sweden | 57°17'05.4''N,<br>14°00'03.9''E | Drainage | Good | 1954 - 2018 | 44 |

### 3.2. Hydrology-related community threshold analysis for peatland taxa

The TITAN analysis of community thresholds for DWT values identified 11 taxa associated with wet conditions (z-) and 21 taxa associated with dry conditions (z+). The change points for individual taxa are presented in Table 2. Among the 32 taxa, 24 were identified as both pure and reliable indicators. The change points for unfiltered data were estimated at 7.4 cm DWT for taxa associated with wetter conditions (Sum(z-) species peak) and 22.1 cm DWT for taxa associated with drier conditions (Sum(z+) species peak) (Fig. 3 & 4). Similarly, the change points derived from the filtered data were 5.8 cm DWT for Sum(z-) and 23.1 cm DWT for Sum(z+), showing only minor differences from the unfiltered data. The range between these values (Sum(z-) 7.4 and Sum(z+) 22.1) represents a potential critical transition zone between wet and dry equilibria, where peatland conditions shift from preserved ecosystems to degraded ones. Among the taxa recognized as both pure and reliable, *Sphagnum papillosum, S. cuspidatum, Andromeda polifolia, S. angustifolium, Rhynospora alba, Scheuchzeria palustris, S. austinii, S. medium/divinum* were assigned to the z-taxa group and taxa such as *S. russowii*, *S. fuscum/rubellum*, *Calluna vulgaris*, Ericaceae, fungal remains of sclerotia, *Erica tetralix*, *S. palustre*, Poaceae, *Hylocomnium splendens*, *Vaccinium oxycoccos*, shrub rootlets, *Empetrum nigrum*, *Pleurozium schreberi*, mycorrhizal roots, *Drosera rotundifolia* and charcoal pieces to the z+ group (taxa in both groups are listed in order of increasing Zenv.cp values; Table 2.).

**Figure. 3.**
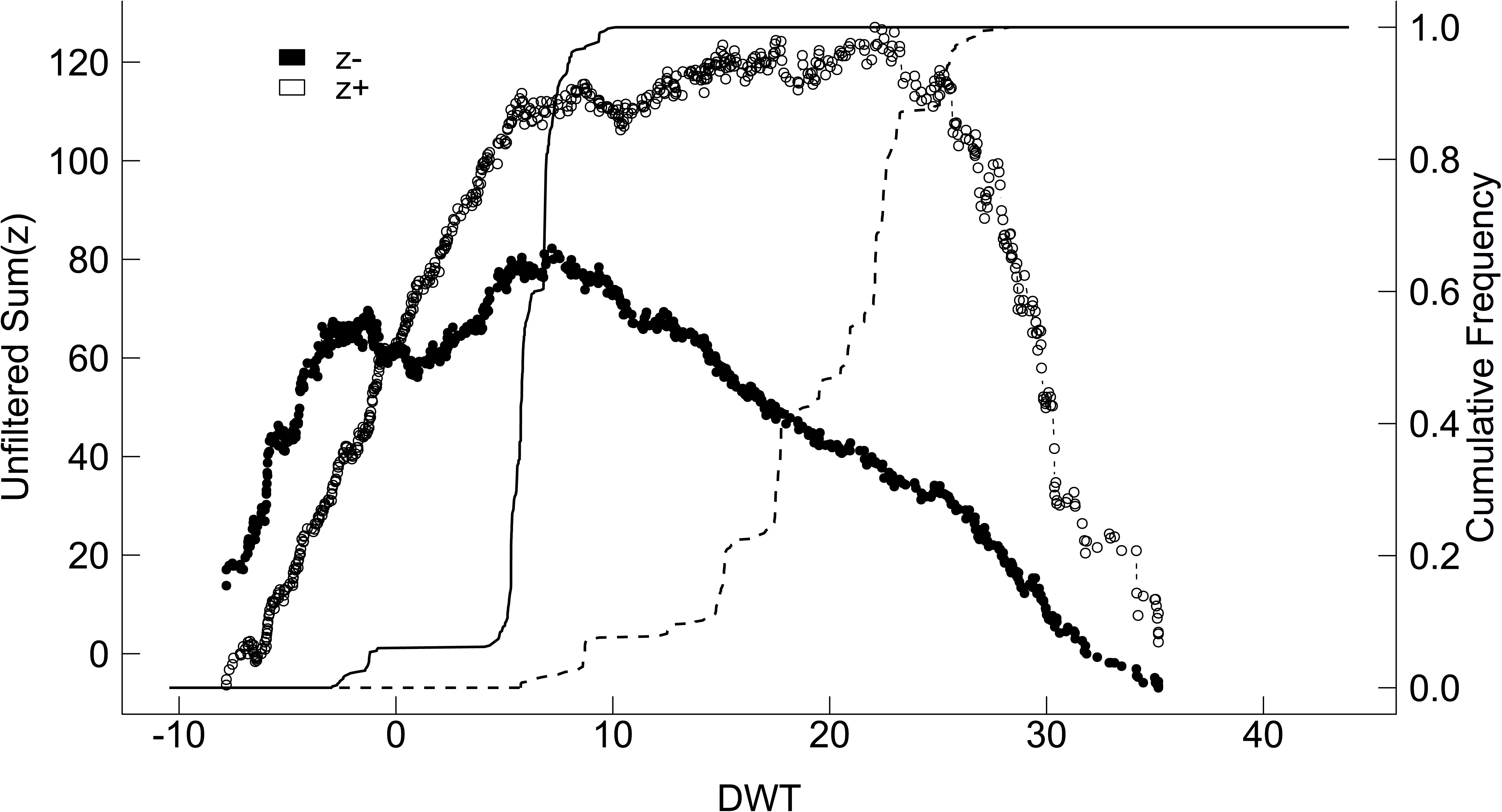
Plot showing TITAN sum(z-) and sum(z+) values for all potential DWT change points. Positive values of the DWT parameter indicate a water table level below the surface, while negative values represent water accumulation above ground level. Peaks in sum(z) indicate the point beyond which synchronous declines in the summarised z-score values occur within the given taxon group. Dot size is proportional to the Z score of a given taxon.

**Table 2.** TITAN-calculated DWT change points for both z- and z+ taxa. Bold text highlights taxa that are both pure and reliable. Zenv.cp – DWT change point based on the Z score; Ienv.cp – DWT change point based on the indicator value score; Z score – Z score value for an individual species; IndVal – indicator value score for an individual species; Frequency – number of subsamples with the species present; Purity – proportion of bootstrap replicates assigning a taxon to the same category; Reliability – proportion of bootstrap replicates with a p-value <0.05 for a given taxon, indicating a predictable response to the DWT gradient.

| Name of the taxon | Zenv.cp | Ienv.cp | Z score | IndVal | Frequency | Purity | Reliability |
| --- | --- | --- | --- | --- | --- | --- | --- |
| **z- taxa** |  |  |  |  |  |  |  |
| *Sphagnum papillosum* | **-2,42** | **-2,51** | **21,88** | **36,14** | **81** | **1,000** | **1,000** |
| *Sphagnum cuspidatum* | **-2,12** | **-7,81** | **28,30** | **57,11** | **160** | **1,000** | **1,000** |
| *Andromeda polifolia* | **-1,44** | **-6,83** | **8,71** | **11,19** | **40** | **0,976** | **0,998** |
| *Sphagnum angustifolium* | **3,62** | **3,52** | **11,17** | **11,39** | **36** | **1,000** | **1,000** |
| *Warnstorfia fluitans* | 5,18 | 5,15 | 4,58 | 2,46 | 7 | 1,000 | 0,926 |
| *Rhynospora alba* | **5,25** | **-3,43** | **5,86** | **6,73** | **35** | **1,000** | **1,000** |
| *Carex limosa* | 7,07 | 6,87 | 2,02 | 0,95 | 3 | 0,928 | 0,400 |
| *Scheuchzeria palustris* | 7,23 | 6,87 | 10,47 | 9,37 | **31** | **1,000** | **1,000** |
| *Pinus mugo* | 12,51 | 16,37 | 2,32 | 2,04 | 10 | 0,868 | 0,780 |
| *Sphagnum austinii* | **12,80** | **12,51** | **8,73** | **14,50** | **76** | **1,000** | **1,000** |
| *Sphagnum medium/divinum* | **17,50** | **25,32** | **17,14** | **58,65** | **433** | **1,000** | **1,000** |
| **z+ taxa** |  |  |  |  |  |  |  |
| *Trichophorum/Eriophorum* | -6,49 | -35,16 | 3,43 | 48,29 | 311 | 0,836 | 0,990 |
| *Sphagnum russowii* | **2,25** | **19,08** | **7,87** | **10,08** | **47** | **1,000** | **1,000** |
| *Sphagnum fuscum/rubellum* | **6,24** | **35,01** | **20,85** | **43,30** | **217** | **1,000** | **1,000** |
| *Calluna vulgaris* | **6,82** | **-7,61** | **8,47** | **8,90** | **37** | **0,976** | **1,000** |
| *Ericaceae* | **8,29** | **6,19** | **19,34** | **26,59** | **99** | **1,000** | **1,000** |
| *Drepanocladus* sp. | 12,49 | 23,23 | 4,62 | 2,16 | 6 | 0,994 | 0,940 |
| **Fungal remains** | **15,03** | **15,18** | **7,15** | **5,34** | **17** | **1,000** | **1,000** |
| *Erica tetralix* | **17,62** | **-7,75** | **9,78** | **7,74** | **20** | **0,972** | **1,000** |
| Herbs | 20,48 | -7,75 | 8,44 | 32,87 | 208 | 0,488 | 1,000 |
| *Sphagnum palustre* | **22,42** | **27,79** | **25,09** | **26,59** | **47** | **1,000** | **1,000** |
| **Poaceae** | **22,62** | **22,86** | **12,37** | **10,17** | **20** | **1,000** | **1,000** |
| *Hylocomnium splendens* | **24,92** | **29,49** | **12,34** | **7,81** | **10** | **1,000** | **1,000** |
| *Oxycoccus* sp. | **25,06** | **27,79** | **10,30** | **15,54** | **53** | **0,998** | **1,000** |
| **Shrubs rootlets** | **25,32** | **26,70** | **10,48** | **54,68** | **371** | **1,000** | **1,000** |
| *Empetrum nigrum* | **26,74** | **26,82** | **12,01** | **6,86** | **7** | **1,000** | **0,994** |
| *Pleurozium schreberi* | **29,10** | **29,76** | **13,20** | **9,56** | **8** | **1,000** | **0,998** |
| *Pinus sylvestris* | 29,10 | -7,82 | 6,63 | 10,81 | 25 | 0,884 | 0,990 |
| **Mycorrhizal roots** | **29,58** | **29,76** | **4,87** | **6,44** | **12** | **0,990** | **0,956** |
| *Drosera rotundifolia* | **30,54** | **31,45** | **11,25** | **12,19** | **6** | **1,000** | **1,000** |
| **Charcoal pieces** | **32,92** | **35,09** | **9,93** | **22,56** | **23** | **0,972** | **1,000** |
| *Sphagnum* sect. *acutifolia* | 34,19 | 34,19 | 4,84 | 12,21 | 10 | 0,990 | 0,928 |

### 3.2 Time-related community threshold analysis for peatland taxa

The TITAN threshold analysis for the presence of peatland taxa over time, spanning from the year 1500 to the present, identified 15 taxa associated with older periods (z-) and 11 taxa linked to more modern timeframes (z+). Among these, seven taxa from each group were classified as both pure and reliable. The change points for individual taxa are provided in Table 3. The community change points for each group in the unfiltered dataset were identified at 1793 for Sum(z−) and 2014 for Sum(z+), respectively (Fig. 4 & 5), and at 1793 and 2013 for the filtered dataset. The interval between these values represents a transition zone between historically dominant plant taxa communities and those currently observed. Distinct peaks for both Sum(z-) and Sum(z+) underscore the statistical significance of these thresholds. Among the taxa identified as both pure and reliable, *Trichophorum/Eriophorum*, *Warnstorfia fluitans*, *Scheuchzeria palustris*, *Sphagnum cuspidatum*, *Rhynospora alba*, *S. medium/divinum* and charcoal pieces were placed within z-group and the following taxa in z+ group: Poaceae, *S. papillosum*, *Erica tetralix*, Ericaceae, *Vaccinium oxycoccos* and *S. palustre* (taxa in both groups are listed in order of increasing Zenv.cp values.)(Table 3.).

**Figure. 4.**
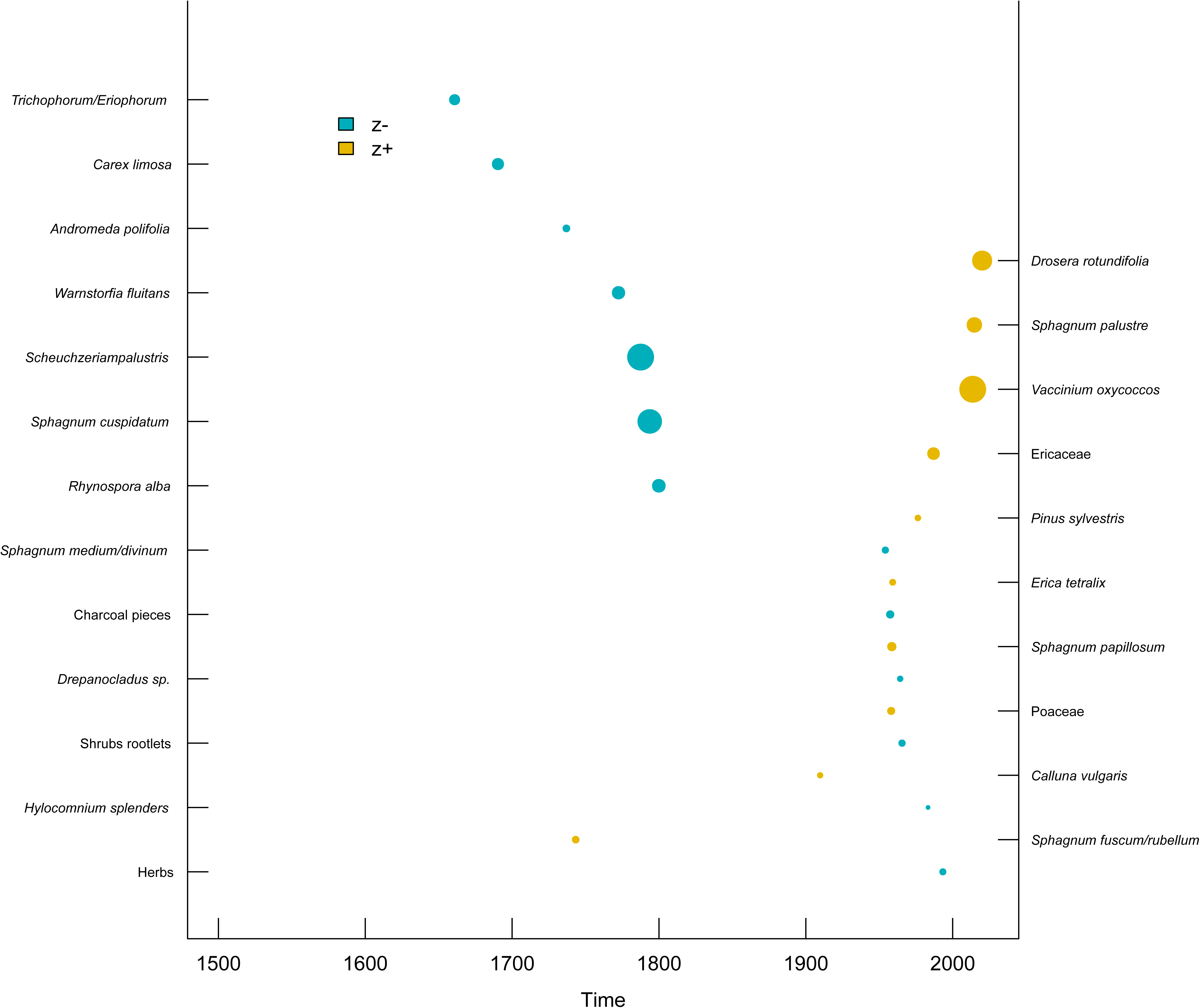
Plot showing taxon-specific change points along the time gradient for taxa predominantly recorded in earlier (z−) versus more recent (z+) periods.

**Figure. 5.**
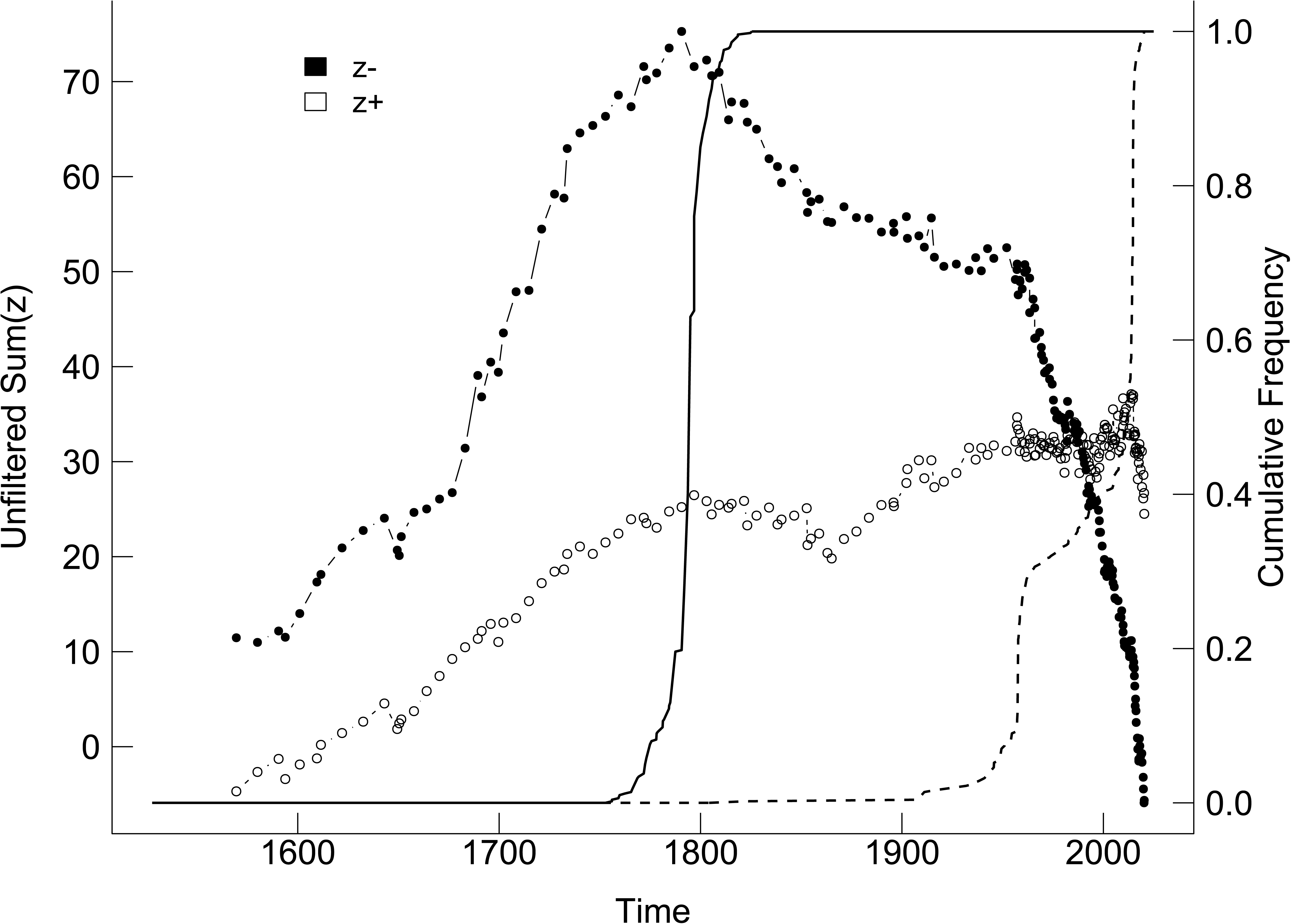
Plot showing TITAN sum(z-) and sum(z+) values for all potential time change points. Peaks in sum(z) indicate the point beyond which synchronous declines in the summarised z-score values occur within the given taxon group. Dot size is proportional to the Z score of a given taxon.

**Table 3.** TITAN-calculated time change points for both z- and z+ taxa. Bold text highlights taxa that are both pure and reliable. Zenv.cp – time change point based on the Z score; Ienv.cp – time change point based on the indicator value score; Z score – Z score value for an individual species; IndVal – indicator value score for an individual species; Purity – proportion of bootstrap replicates assigning a taxon to the same category; Reliability – proportion of bootstrap replicates with a p-value <0.05 for a given taxon, indicating a predictable response to the time gradient.

| **Name of the taxon** | **Zenv.cp** | **Ienv.cp** | **Z score** | **IndVal** | **Frequency** | **Purity** | **Reliability** |
| --- | --- | --- | --- | --- | --- | --- | --- |
| **z- taxons** |  |  |  |  |  |  |  |
| *Trichophorum/Eriophorum* | **1660,85** | **1646,2** | **8,12** | **69,64** | **67** | **1,000** | **0,998** |
| *Carex limosa* | 1690,35 | 1690,35 | 9,14 | 13,04 | 3 | 0,946 | 0,784 |
| *Andromeda polifolia* | 1736,95 | 1610,50 | 4,93 | 11,04 | 6 | 0,906 | 0,910 |
| *Warnstorfia fluitans* | **1772,45** | **1736,95** | **10,24** | **17,95** | **7** | **1,000** | **0,998** |
| *Scheuchzeria palustris* | **1787,50** | **1793,70** | **23,27** | **71,28** | **31** | **1,000** | **1,000** |
| *Sphagnum cuspidatum* | **1793,70** | **1781,30** | **21,04** | **60,05** | **30** | **1,000** | **1,000** |
| *Rhynospora alba* | **1799,90** | **1574,75** | **10,56** | **27,21** | **17** | **1,000** | **1,000** |
| *Sphagnum medium/divinum* | **1954,15** | **1616,75** | **4,42** | **42,46** | **116** | **0,976** | **0,994** |
| **Charcoal pieces** | **1957,45** | **1957,45** | **5,40** | **9,41** | **8** | **1,000** | **0,982** |
| *Drepanocladus* sp. | 1964,25 | 1963,40 | 3,63 | 6,38 | 6 | 0,998 | 0,832 |
| Shrubs rootlets | 1965,50 | 1605,25 | 4,50 | 48,34 | 115 | 0,840 | 1,000 |
| *Hylocomnium splendens* | 1983,20 | 1982,70 | 2,10 | 5,04 | 6 | 0,914 | 0,604 |
| *Pleurozium schreberi* | 1992,70 | 1992,7 | 2,09 | 6,06 | 8 | 0,844 | 0,628 |
| Herbs | 1993,30 | 2016,10 | 4,31 | 34,63 | 73 | 0,786 | 0,998 |
| *Empetrum nigrum* | 1998,30 | 1998,30 | 1,35 | 4,90 | 7 | 0,730 | 0,450 |
| **z+ taxons** |  |  |  |  |  |  |  |
| *Sphagnum fuscum/rubellum* | 1743,30 | 2019,15 | 3,97 | 46,30 | 100 | 0,796 | 1,000 |
| Mycorrhizal roots | 1843359 | 1852,90 | 1,85 | 6,12 | 9 | 0,744 | 0,594 |
| *Calluna.vulgaris* | 1909,75 | 1697,70 | 2,90 | 14,27 | 27 | 0,962 | 0,918 |
| **Poaceae** | **1958,10** | **2014,75** | **4,30** | **13,49** | **20** | **1,000** | **0,982** |
| *Sphagnum papillosum* | **1958,55** | **1958,55** | **5,24** | **13,84** | **17** | **0,988** | **1,000** |
| *Erica tetralix* | **1959,15** | **1959,15** | **3,37** | **12,08** | **19** | **0,962** | **0,932** |
| *Pinus sylvestris* | 1976,30 | 2018,00 | 3,11 | 10,05 | 16 | 0,924 | 0,880 |
| **Ericaceae** | **1986,95** | **1986,95** | **7,91** | **28,41** | **39** | **1,000** | **1,000** |
| *Oxycoccus* sp. | **2013,65** | **2017,75** | **19,15** | **81,28** | **45** | **1,000** | **1,000** |
| *Sphagnum palustre* | **2014,75** | **2020,35** | **10,14** | **49,94** | **42** | **1,000** | **1,000** |
| *Drosera rotundifolia* | 2020,05 | 2020,35 | 13,72 | 56,14 | 6 | 0,984 | 0,928 |

### 3.3 Effect of changing DWT on the plant taxonomic composition in peat cores

The analysis of plant macrofossil composition found in peat cores, combined with testate amoebae-derived DWT values, revealed significant differences in the hydrological histories of the seven studied peatlands (Fig. 6). All sites experienced substantial fluctuations in water levels, with some changes particularly prominent. Among them, only Bagno Kusowo exhibited an improvement in DWT value in the second half of the 20th century, with DWT increasing from approximately 20 cm to around 12 cm below the surface. This shift was accompanied by a transition in the dominant *Sphagnum* species, from *S. fuscum/rubellum* (classified in the first TITAN analysis as a drought-adapted taxon from the z+ group, Table 2) to *S. medium/divinum* (associated with the more water-dependent z− group).

**Figure. 6.**
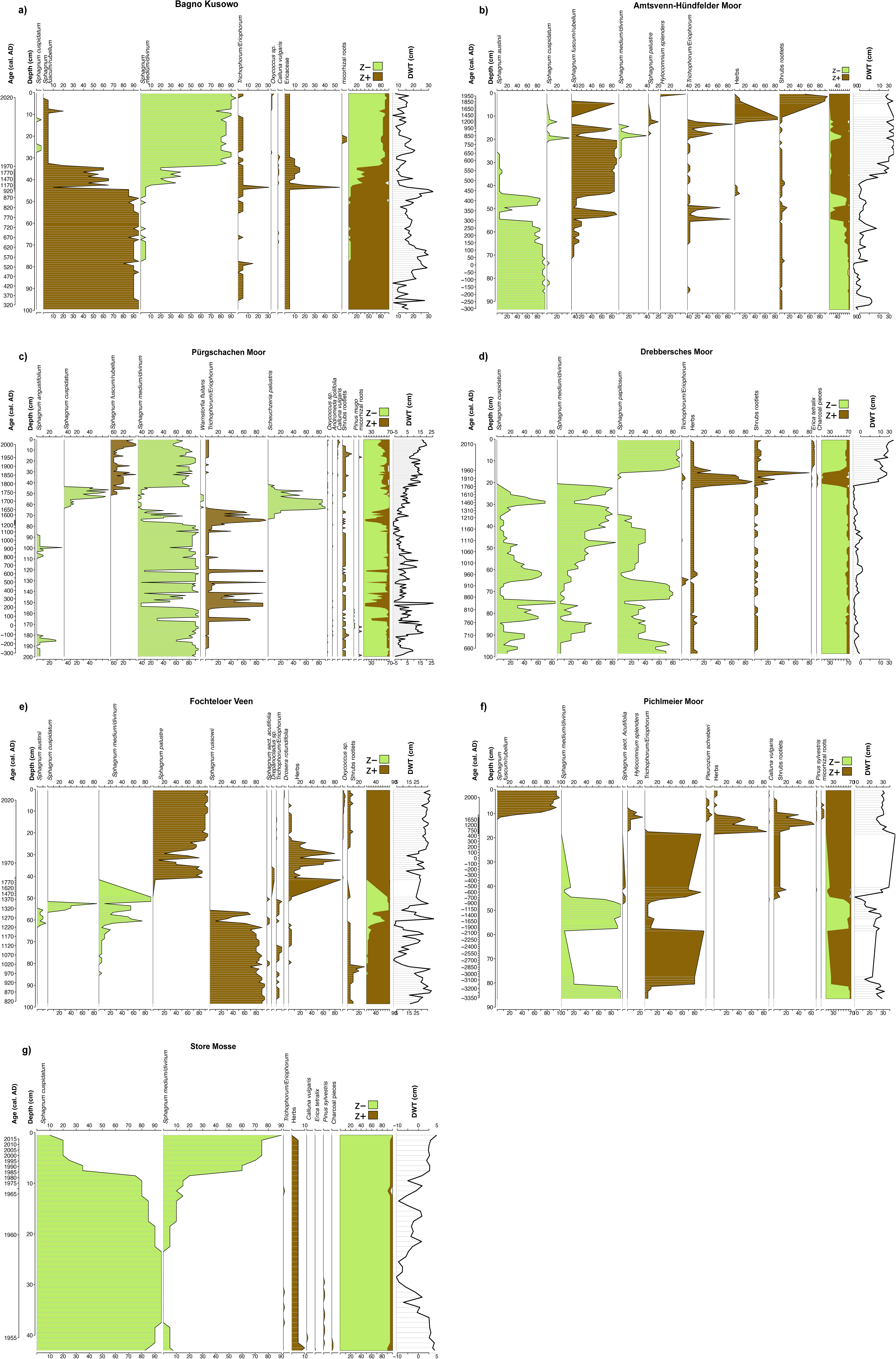
Diagrams showing the proportion of dominant macrofossil taxa in the analysed peat core layers, along with estimated water level changes and shifts in the ratios of z+ (wet associated taxa - green) and z-species (dry associated taxa - brown). Negative values of the DWT parameter indicate a water table level below the surface, while positive values represent water accumulation above ground level. The full set of the age-depth models is presented in the Supplementary Material (Supplementary Figure 1-7), complete plant macrofossils diagrams are presented in the Supplementary Material (Supplementary Figure 8-14), and complete testate amoebae diagrams are presented in the Supplementary Material (Supplementary Figure 15-21).

An especially notable and abrupt decline in the water table, leading to significant shifts in dominant plant taxa, was observed at Amtsvenn-Hündfelder Moor and Drebbersches Moor. In these sites, the DWT values changed from around 5 cm to 25 cm and from −5 cm to 25 cm, respectively. At Amtsvenn-Hündfelder Moor, *S. austinii* (z− group), which thrived under more favourable hydrological conditions, disappeared, being replaced by *S. fuscum/rubellum* (z+ group) along with an increased presence of shrub rootlets (z+). A similar pattern was recorded at Drebbersches Moor, where declining water levels led to the replacement of *S. cuspidatum* and *S. medium/divinum* (both z−taxa) by *S. papillosum* and the emergence of vascular plants such as *Erica tetralix* (z+), as well as herbs (z+) and additional shrubs (z+), particularly over the last few decades.

Despite covering only the last few decades, the core of Store Mosse also revealed an intriguing shift in dominant *Sphagnum* species. For several years, *S. cuspidatum* (z−) was the dominant species in the analysed profile. However, following a drop in DWT around 1990 (to ∼4 cm), which persisted in subsequent years, *S. medium/divinum* (z−) gradually replaced it. Although both species were allocated to the z− group, their estimated DWT change points differ strongly (*S. cuspidatum*: −2.12 cm, *S. medium/divinum*: 17.50 cm), indicating a shift within the z− group toward a species less dependent on high moisture levels. A similar transition within the same z group was observed at Fochteloer Veen. Here, DWT fluctuations over centuries led to the dominance of *S. russowii* (z+), which was interrupted by a two-century-long period of *S. medium/divinum* (z−) dominance, during which the estimated DWT rose by a few centimetres. After 1950, the DWT value dropped and stabilised at a low level of around 25 cm, which was followed by the dominance of *S. palustre* (z+) and a noticeable increase in vascular plant presence, including *Vaccinium oxycoccos* (z+) and *Drosera rotundifolia* (z+). Although both *S. russowii* and *S. palustre* belong to the z+ group, the transition from a species with a higher to a lower DWT change point (2.25 cm to 22.42 cm, respectively) suggests a long-term deterioration of hydrological conditions in recent decades.

Pichlmeier Moor has undergone substantial hydrological changes over the past centuries. Initially characterised by a relatively low water table (DWT ∼24 cm) and dominance of *S. medium/divinum* (z−), the site experienced significant drying (DWT ∼36 cm), leading to the significant development of vascular taxa such as *Trichophorum/Eriophorum*. However, in the last few decades, a relative improvement in hydrological conditions (DWT ∼28 cm) has led to the re-establishment of *Sphagnum* dominance, specifically *S. fuscum/rubellum* (z+).

At Pürgschachen Moor, the early centuries were characterised by consistently low DWT values (mostly below 10 cm) and the dominance of *S. medium/divinum* (z−, estimated DWT change point: 15.53 cm). Around the 17th century, the site experienced a temporary increase in the water table (DWT ∼−5 cm), coinciding with a shift in dominant *Sphagnum* species to *S. cuspidatum* (z−, estimated DWT change point: −2.12 cm). Over the last two centuries, a continuous decline in hydrological conditions (DWT reaching ∼15 cm) has been observed, marked by the resurgence of *S. medium/divinum* (z−) and the increasing dominance of *S. fuscum/rubellum* (z+).

## 4. Discussion

### 4.1 Critical Thresholds of water level in ombrotrophic peatlands

Appropriate water levels, measured as DWT, remain a key parameter for the conservation and maintenance of peatlands in a favourable ecological state (Haapalehto et al. 2011, Waddington et al. 2015, Kreyling et al. 2021, Baird and Low 2022, Zak and McInnes 2022). Our research identified two change points for both taxa that benefit from lower and higher DWT levels. These findings indicate a transition zone between the dominance of species classified as moisture-dependent and those more drought-adapted, occurring between 7.2 and 22.1 cm DWT. According to our results, a decline in DWT within this range may promote a shift in plant species composition toward more drought-adapted taxa, with the likelihood of such a transition increasing as the water level drops even further.

The defined critical transition zone is broader yet consistent with previous estimations by Lamentowicz et al. (2019), who identified a transition range of 8 to 17 cm, with a tipping point at 11.69 cm using nearly identical methods. Similarly, the guidelines provided by Couwenberg (2011) suggest an optimal DWT of approximately ±10 cm, aligning with values reported to support the restoration of previously degraded peatlands (Haapalehto et al. 2011). In contrast, Jassey et al. (2018), based on short-term experimental manipulations (2 years) and subsequent shifts in both plant and fungal communities, estimated the transition zone to lie between 19 and 23 cm DWT - close to our estimated tipping point for z+ species. Additionally, long-term analyses of peat accumulation rates across several European peatlands have shown that the optimal DWT level for the highest peat accumulation lies between 5 and 10 cm (Swindles et al. 2025). In summary, based on both our findings and previous research, we propose that the critical transition zone for plant communities ranges from 15 to 22 cm, above which a rapid species shift may occur. Meanwhile, a DWT level of 10 cm could be recommended as optimal and safe from the perspective of peatland restoration and conservation.

Empirical evidence supporting these values is also found in macrofossil analyses. For instance, records from Drebbersches Moor and Amtsvenn-Hündfelder Moor document a rapid turnover in plant species composition in response to declining DWT levels. Both sites were initially dominated by *Sphagnum* species from the z-group (such as: *S. cuspidatum, S. austinii* or *S. medium/divinum*) until the DWT exceeded 20 cm, leading to a significant shift in dominant moss species to those from z+ group (*S. fuscum/rubellum*) as well as the introduction of a higher proportion of vascular plant species (*Erica tetralix* and shrubs). Another case supporting the estimated values, where an increase in water level promoted a shift in the opposite direction, was observed in the partially restored Bagno Kusowo peatland. At this site, hydrological improvements led to a DWT change from ∼20 cm to 12 cm. Following this shift, a turnover in the dominant *Sphagnum* taxa occurred, with the more drought-adapted *S. fuscum* (z+ taxon) being replaced by *S. medium/divinum* (z-taxon). While these observations are based on individual case studies, they support the estimations from our research and previous studies.

It is essential to note the similarity between the results of our analysis and those of Lamentowicz et al. (2019), who, for their DWT tipping point estimation, used almost identical methods. However, in their work, the data - although originating from multiple and diverse peatlands - came from only one European country, raising the question of whether these results could be applied to peatlands at the continental or even global level. The fact that our study, which utilized the same TITAN method, but with data from seven distinct sites across the European continent (with only one site shared with Lamentowicz et al. 2019, for which we present original and independent data), yielded similar results further strengthens the reliability of the estimated community thresholds. In addition, it is worth noting that the peatlands examined in our study differ somewhat from those of Lamentowicz et al. (2019), as all of our sites have experienced significant historical drainage and peat excavation and are currently undergoing a restoration process.

It should be emphasised that a species shift due to a declining water level does not necessarily indicate a loss of peatland functionality or ecological value. A comprehensive analysis of 56 European peatland sites by Robroek et al. (2017) examined species composition across different environmental gradients and found that while climatic factors may drive species turnover, the overall functional composition of these ecosystems often remains largely unchanged. This can be explained by the fact that disappearing species are replaced by taxa that, although better adapted to new environmental conditions, still fulfil similar ecological roles. These findings suggest that peatlands exhibit a degree of resilience to environmental changes, including those driven by climate change. Therefore, peatlands that have undergone significant species turnover in the past should still be considered vital ecosystems, with their continued conservation remaining a priority.

### 4.2 Time

Currently, a significant portion of the world’s peatlands is classified as degraded. In countries historically rich in peatlands, such as Germany, the Netherlands, Poland or Finland, a substantial proportion of these ecosystems has been lost or severely altered (Tanneberger et al. 2021a). Our TITAN analyses, combined with peat macrofossil and DWT reconstructions, provide valuable insights into the history of peatland degradation across Europe. The findings suggest a pronounced shift in taxonomic composition over time, particularly during the 19th and 20th centuries, with a marked acceleration after 1950-likely in response to increasing ecosystem degradation. This interpretation aligns with historical records (Lamentowicz et al. 2015), which indicate that extensive peat extraction took place from the late 18th century through the first half of the 20th century, typically following drainage efforts. Furthermore, testate amoeba data from multiple European sites suggest that the period between 1800 and 2000 was the driest in the long-term history of these ecosystems (Swindles et al. 2019), reinforcing the hypothesis that significant shifts in species composition occurred during this time. Following World War II (1939-1945 CE), the exploitation, drainage, and degradation of peatlands intensified across several European countries, leading to substantial declines in water table levels (Swindles et al. 2019). In some cases, these processes were so intensive that, for example, in Finland, estimates suggest that up to 70% of the country’s mires were lost due to ambitious drainage projects, which reached their peak in the 1970s (Tanneberger et al. 2021a). This large-scale decrease in water level may be reflected in our second TITAN analysis, which shows a sharp decline in sum(z−) species and an exceptionally rapid increase in sum(z+) species after 1950. Furthermore, the analysis identified a notable shift towards vascular plants, particularly drought-adapted taxa such as *Vaccinium oxycoccos*, Ericaceae, *Pinus sylvestris*, Poaceae, and *Drosera rotundifolia*. Similarly, *Sphagnum* taxa (*S. palustre*, *S. papillosum*, and *S. fuscum/rubellum*), while still present in z+ group, included two species that were classified in the first TITAN analysis as part of the z+ group (associated with lower DWT conditions), indicating a shift toward more drought-adapted *Sphagnum* species.

Overall, our findings suggest that not only has a significant turnover in species composition occurred over the past 200 years, with an acceleration in recent decades, but also that this shift reflects a long-term decline in water availability (Giese et al. 2025). The dominance of drought-adapted species in degraded peatlands supports the hypothesis that hydrological conditions have progressively deteriorated, particularly in the latter half of the 20th century. This aligns well with global wetland change reconstructions (Fluet-Chouinard et al. 2023), which indicate that since 1700, nearly 50% of northern European wetlands, including peatlands, have been lost, with a significant portion disappearing in the second half of the 20th century.

### 4.3 Implications for restoration

In recent years, increasing attention has been paid to the risks associated with leaving degraded peatlands unmanaged (Günther et al. 2020, Tanneberger et al. 2020, 2021b, Zak and McInnes 2022). Globally, it is estimated that even 15% of the total peatland area has been drained (Joosten and Clarke 2002) with such sites potentially accounting for as much as 5% of global anthropogenic COLJ emissions (Joosten and Clarke 2002, Friedlingstein et al. 2019), not to mention emission of the other important greenhouse gases such as CH_4_ or N_2_O. Emissions from these degraded ecosystems represent a substantial challenge in the fight against climate change, particularly in Europe, which is the biggest greenhouse gas (GHG) emitter from degraded peatlands just after Indonesia (Joosten 2010). Historically, the continent was extensively covered by peatlands, but it is now estimated that approximately 25% of Europe’s peatlands remain degraded, with this figure rising to as much as 50% within the European Union (Tanneberger et al. 2021a). The costs of leaving peatlands in an unsatisfactory state are therefore considerable, making their restoration a crucial task for reducing GHG emissions.

Fortunately, research and practical experience demonstrate that peatland restoration is feasible (Zak and McInnes 2022). While the initial stages of rewetting of degraded peatlands are associated with substantial methane emissions, rewetting has been shown to rapidly and significantly reduce COLJ emissions (Huth et al. 2022). Although methane’s impact on global warming is considerable, as Günther et al. (2020) highlighted in their work, its atmospheric lifespan is much shorter than that of COLJ (approximately 12 years compared to 100 years). Therefore, in the long term, rewetting a degraded peatland is always a more favourable solution than leaving it unmanaged, as it increases the amount of C accumulated in the peat (Günther et al. 2020, Tanneberger et al. 2021b). Moreover, studies suggest that methane emissions during the rewetting process can be mitigated. For example, the removal of the degraded top layer of peat during topsoil removal has been shown to significantly reduce methane emissions (Zak and McInnes 2022). In addition, it should be noted that both conservation and restoration of peatlands support most of the United Nations Sustainable Development Goals, for example, such as “no poverty”, “clean water and sanitation”, “climate action”, or “life on land” (Tanneberger et al. 2020). Thus, while restoring proper hydrological conditions is clearly a priority, the question remains as to what level of DWT is optimal for facilitating the regeneration of degraded peatlands and ensuring the long-term preservation of still-intact peatlands.

Based on our palaeoecological findings and available data, we propose that a safe DWT for peatland regeneration is approximately 10 cm below the surface. Negative deviations from this value are associated with an increased likelihood of shifts in species composition towards drought-adapted taxa. Accordingly, our results suggest that the DWT dropping to the range of ∼ 15 to 22 cm increases the risk of a significant species turnover, thus marking this as a transition zone towards a more drought-resistant species. While these values offer essential insights into the relationship between water levels and plant communities, their application requires caution. Different peatland sites may exhibit varying degrees of tolerance to drainage, leading to site-specific differences in their critical DWT thresholds. Despite these variations, the estimated values remain highly relevant for guiding practical conservation and management efforts to maintain peatland ecosystems in a healthy state.

Furthermore, although this proposed DWT level may appear stringent - particularly given that in many degraded peatlands, the water table is often several dozen centimeters below the surface, it should not discourage restoration efforts, even if the target DWT of 10 cm could not be achieved immediately, nor should it detract from the importance of protecting such areas. Primarily, even the peatlands that show signs of degradation and have already undergone a species transition can still fulfil several crucial ecological roles. Studies show that while a species shift is undoubtedly a momentous event that has no small meaning for biodiversity protection, in practice, vanishing species are often replaced by new species which are not only more tolerant to the novel conditions but often also fulfil similar ecological functions (Robroek et al. 2017). This information is crucial as it also stresses the purpose of protecting peatlands, even those that underwent essential changes over the years. This situation is also visible up to a degree in our data, for example, key *Sphagnum* taxa are present in both z+ and z-groups. At the same time, the DWT threshold for different species differs significantly.

What is worth noting is that the DWT level of around 10 cm may also be an essential threshold from the perspective of mitigating carbon emissions from peatlands. Such conclusions regarding carbon emissions were made by Evans et al. (2021), who analysed the relationship between peatland DWT levels and carbon fluxes using the data collected from multiple eddy covariance flux towers located near English and Irish peatlands. Their study revealed that higher DWT levels were associated with substantial COLJ emissions, whereas lower DWT levels promoted increased CHLJ emissions. However, what is worth noticing is that a DWT level of approximately 10 cm facilitated net COLJ uptake from the atmosphere, effectively offsetting the CHLJ emissions observed at this DWT level. This balance should result in an overall absorption of greenhouse gases, highlighting the potential importance of maintaining such low DWT levels to mitigate climate change. These findings are consistent with the observations of Fortuniak et al. (2021), who, based on six years of continuous eddy covariance COLJ and CHLJ flux measurements from a single site, concluded that to mitigate greenhouse gas emissions, the DWT should not fall below 15 cm. Similarly, Chimner et al. (2017), in their analysis of the long-term effects of water level manipulation on GHG emissions, reported that a site with a DWT of approximately 12 cm acted as the highest carbon sink compared to areas with lower water levels. Thus, both these studies and our own results suggest that a DWT of around 10 cm may be the most optimal for both peatland preservation and carbon emissions mitigation. Such a water level is likely to not only preserve the ecosystem’s integrity or stimulate the regeneration of degraded peatlands but also, within a relatively short period, contribute to efforts to slow the growth of GHG concentration in the atmosphere. This is particularly important, as many drained peatlands currently act as significant carbon emitters (Leifeld et al. 2019, Doelman et al. 2023).

It is crucial to emphasise that, while peatland restoration is achievable, it is neither a straightforward process nor a guaranteed success, as it heavily depends on a thorough understanding of the local hydrological regime (Zak and McInnes 2022). Therefore, priority should be given to protecting peatlands that remain in good condition before they degrade. Our analysis of peat cores indicates that even a relatively short time of water regime disruption can result in significant shifts in peatland species composition. This rapid response to hydrological changes has also been documented in other studies (Couwenberg et al. 2011). In contrast, practical experience demonstrates that peatland restoration is a much more prolonged and complex endeavour, often taking decades to achieve meaningful results (Kreyling et al. 2021). Restoration efforts are typically resource-intensive and costly, while the degradation of a peatland can occur within just a few years of unfavourable conditions. Furthermore, even successfully restored peatlands often exhibit significant differences in biodiversity and species composition compared to those that have never been degraded (Kreyling et al. 2021). Thus, while the functionality of peatland may be restored up to a degree, its pristine character may never be recovered in a matter of a few decades (Haapalehto et al. 2011). This issue is also evident in the case of microbial data from Bagno Kusowo. Although the site has undergone a self-rewetting process in recent decades, its microbial community remains significantly different from the typical species composition observed in similar but non-degraded part of the same site. Consequently, while restoration is achievable, it is a challenging and time-consuming process, highlighting the importance of proactive protection measures to prevent degradation in the first place.

Unfortunately, only a limited proportion of peatlands are currently subject to any form of formal protection, and even among those that are protected, many face inadequate safeguards (Tanneberger et al. 2021a). For instance, in some cases, peatlands are included within the boundaries of protected areas, but they are not the primary focus of conservation efforts. In other instances, the protection is confined to the peatland body itself, leaving the adjacent drainage basin unprotected, which may threaten the hydrological stability of the area, essential for peatland preservation. Therefore, the authors of this study call for addressing these gaps in conservation strategy and applying proper rewetting strategies as a vital step to ensuring the long-term integrity and functionality of these ecosystems.

### Conclusions

Our integrated analysis of testate amoeba-derived DWT reconstructions and plant macrofossil records identified a critical transition zone between 7 and 22 cm below the peat surface. Crossing this threshold is associated with significant shifts in plant species composition, including a decline in *Sphagnum* taxa, which likely signals the onset of peatland degradation. Microbial community data from Bagno Kusowo further reveal that, even following hydrological restoration, microbial assemblages differ markedly from those found in undisturbed, pristine peatlands. This suggests that biological recovery may lag behind hydrological improvements or follow a different trajectory altogether. Moreover, analysis of vegetation dynamics across the studied sites indicates a pronounced shift from moisture-dependent to drought-adapted plant species beginning in the early 18th century and accelerating in the latter half of the 20th century-corresponding with increased anthropogenic pressure and a widely documented drop in water level. Based on our multi-proxy data and long-term ecological trajectories, as well as evidence from previous studies, we recommend maintaining a mean DWT of approximately 10 cm below the surface as an optimal target for both peatland protection and restoration efforts.

## Acknowledgements

This research was funded through the 2020–2021 Biodiversa+ and Water JPI joint call for research projects, under the BiodivRestore ERA-NET Cofund (GA N◦ 101003777), with the EU and the funding organisations National Science Centre (Poland) proposal number - 2021/03/Y/ST10/00093 (Poland), DFG (Germany), FWF (Austria), and the Ministry of LNV (The Netherlands). The authors are grateful to Patryk Fiutek for the help in the field and for analysing testate amoebae.

## Abbreviations

TITAN: threshold indicator taxa analysis
DWT: depth to water table

