## Supplementary materials Captions for "Long-Term Ecological Baselines and Critical Thresholds in Ombrotrophic Peatlands of Europe: Implications for Restoration Strategies"

**Supplementary Material**

**Supplementary Figures**

- **Supplementary Figure 1** — Bayesian age-depth model (based on ¹⁴C dating), *Amtsvenn-Hündfelder Moor*
- **Supplementary Figure 2** — Bayesian age-depth model (based on ¹⁴C dating), *Bagno Kusowo*
- **Supplementary Figure 3** — Bayesian age-depth model (based on ¹⁴C dating), *Drebbersches Moor*
- **Supplementary Figure 4** — Bayesian age-depth model (based on ¹⁴C dating), *Fochteloer Veen*
- **Supplementary Figure 5** — Bayesian age-depth model (based on ¹⁴C dating), *Pichlmeier Moor*
- **Supplementary Figure 6** — Bayesian age-depth model (based on ¹⁴C dating), *Pürgschachen Moor*
- **Supplementary Figure 7** — Bayesian age-depth model (based on ¹⁴C dating), *Store Mosse*
- **Supplementary Figure 8** — Plant macrofossils diagram, *Amtsvenn-Hündfelder Moor*
- **Supplementary Figure 9** — Plant macrofossils diagram, *Bagno Kusowo*
- **Supplementary Figure 10** — Plant macrofossils diagram, *Drebbersches Moor*
- **Supplementary Figure 11** — Plant macrofossils diagram, *Fochteloer Veen*
- **Supplementary Figure 12** — Plant macrofossils diagram, *Pichlmeier Moor*
- **Supplementary Figure 13** — Plant macrofossils diagram, *Pürgschachen Moor*
- **Supplementary Figure 14** — Plant macrofossils diagram, *Store Mosse*
- **Supplementary Figure 15** — Percentage testate amoebae diagram, *Amtsvenn-Hündfelder Moor*
- **Supplementary Figure 16** — Percentage testate amoebae diagram, *Bagno Kusowo*
- **Supplementary Figure 17** — Percentage testate amoebae diagram, *Drebbersches Moor*
- **Supplementary Figure 18** — Percentage testate amoebae diagram, *Fochteloer Veen*
- **Supplementary Figure 19** — Percentage testate amoebae diagram, *Pichlmeier Moor*
- **Supplementary Figure 20** — Percentage testate amoebae diagram, *Pürgschachen Moor*
- **Supplementary Figure 21** — Percentage testate amoebae diagram, *Store Mosse*

**Supplementary Tables**

- **Supplementary Table 1** — The list of radiocarbon dates from seven studied peatlands with calibration. The IntCal20 (Reimer et al., 2020) and Bomb21NH1 (Hua et al., 2021) atmospheric curves were used to calibrate the dates. *pMC*– percent modern carbon.
