## Supplemental Data 1 for "Long-Term Ecological Baselines and Critical Thresholds in Ombrotrophic Peatlands of Europe: Implications for Restoration Strategies": Tab. 1. Radiocarbon dates. various cores.docx

Tab. 1. Radiocarbon dates from various sites. Biodiversa project

| **Depth (cm)** | **Material** | **Nr. Lab.** | **C14 date** | **Age cal yr BP (95.4 %)** | **Age BC/AD** |
| --- | --- | --- | --- | --- | --- |
|  | **POLAND, Bagno Kusowo** | | |  |  |
| KUSREV 15-16 | *Sphagnum* stems | Poz-152219 | 104.53 ± 0.34 pMmCodern |  |  |
| KUSREV 30-31 | *Sphagnum* stems | Poz-152220 | 114.85 ± 0.36 pMmCodern |  |  |
| BK 35.5 | *Sphagnum* stems | Poz-161646 | 135.07 ± 0.39 pMC |  |  |
| KUSREV 45-46 | *Sphagnum* stems | Poz-152626 | 1265 ± 30 BP |  |  |
| KR1 59-60 | *Sphagnum* stems | Poz-156100 | 1450 ± 30 BP |  |  |
| KR1 79-80 | *Sphagnum* stems | Poz-156101 | 1530 ± 30 BP |  |  |
| KR1 98-99 | *Sphagnum* stems | Poz-156103 | 1645 ± 30 BP |  |  |
|  | **GERMANY, Vechtaer Moor** | | |  |  |
| **Depth (cm)** | **Material** | **Nr. Lab.** | **C14 date** | **Age cal yr BP (95.4 %)** | **Age BC/AD** |
| DM1 14-15 | *Sphagnum* stems | Poz-158121 | 105.93 ± 0.33 pMC modern | 255-33 | 1695-1917 |
| DM1 17-18 | *Trichophorum cespitosum* fruits | Poz-158753 | 115.68 ± 0.36 pMC modern | 295-31 | 1692-1919 |
| DM1 35-36 | *Sphagnum* stems*, Carex* sp. fruit, *Rhynospora alba* fruit | Poz-158122 | 900 ± 30 BP | 909-732 | 1042-1219 |
| DM1 73-74 | *Sphagnum* stems | Poz-158123 | 1200 ± 30 BP | 1244-1006 | 706-945 |
| DM1 98-99 | *Sphagnum* stems, *Oxycoccus palustris* leaves*, Andromeda polifolia* leaves | Poz-159301 | 1370 ± 30 BP | 1346-1178 | 605-772 |
|  | **GERMANY, Amtsvenn Moor** | | |  |  |
| AV1 7-8 | Baeothryon *cespitosum* fruits | Poz-158751 | 130 ± 30 BP | 275-8 | 1675-1942 |
| AV1 15-16 | *Sphagnum* stems | Poz-158752 | 1160 ± 30 BP | 1178-973 | 773-978 |
| AV1 27-28 | *Sphagnum* stems | Poz-158118 | 1480 ± 30 BP | 1400-1307 | 550-644 |
| AV1 55-56 | *Sphagnum* stems | Poz-158119 | 1670 ± 30 BP | 1693-1420 | 275-531 |
| AV1 93-95 | *Sphagnum* stems | Poz-158107 | 2095 ± 30 BP | 2146-1949 | -197 - 1 |
|  | **Austria, Purgschachen Moor** | | |  |  |
| PM1 16,5 | *Sphagnum* stems | Poz-158108 | 103.01 ± 0.33 pMC modern |  | 1696-1914 |
| PM1 44,5 | *Sphagnum* stems, *Oxyccocus palustris* leaves, *Rhynospora alba* fruit | Poz-158109 | 145 ± 30 BP |  |  |
| PM1 65,5 | *Sphagnum* stems | Poz-164571 | 170 ± 30 BP |  |  |
| PM1 79,5 | *Sphagnum* stems | Poz-158110 | 950 ± 30 BP |  | 1028-1162 |
| PM1 115,5 | *Sphagnum* stems | Poz-158111 | 1320 ± 30 BP |  | 652-775 |
| PM1 156,5 | *Pinus mugo* cone | Poz-158112 | 1880 ± 30 BP |  | 81-236 |
| PM1 198,5 | *Sphagnum* stems | Poz-158120 | 2165 ± 30 BP |  | -357- 60 |
|  | **Austria, Pilchmeier Moor** | | |  |  |
| PI1 11,5 | *Pleurozium schreberi* stem with leaves + *Hylocomnium splenders* stems with leaves | Poz-159505 | 102.78 ± 0.32 pMC |  |  |
| PI1 19,5 | *Sphagnum* stems | Poz-159506 | 1770 ± 30 BP |  |  |
| PI1 45,5 | *Sphagnum* stems | Poz-159507 | 2450 ± 30 BP |  |  |
| PI1 65,5 | *Sphagnum* stems | Poz-159508 | 4215 ± 35 BP |  |  |
| PI1 85,5 | *Sphagnum* stems | Poz-159509 | 4415 ± 35 BP |  |  |
|  | **Niederlands, Fochtelooër Veen** | | |  |  |
| FV1 16,5 | *Oxycoccus palustris leaves* | Poz-159510 | 101.47 ± 0.34 pMC modern |  | 1697-1911 |
| FV1 40,5 | *Sphagnum* stems | Poz-159303 | 104.08 ± 0.35 pMC modern |  | 1696-1916 |
| FV1 47,5 | *Sphagnum* stems and leaves, charcoal pieces | Poz-159512 | 570 ± 30 BP |  | 1306-1424 |
| FV1 62,5 | *Sphagnum* stems | Poz-159513 | 820 ± 30 BP |  | 1175-1273 |
| FV1 98,5 | *Sphagnum* stems | Poz-159514 | 1215 ± 30 BP |  | 689-890 |
|  | **Stoore Moose, Sweden** | | | |  |
| SM13.5 | *Sphagnum* stems | Poz-172186 | 100.67 ± 0.34 pMC |  |  |
| SM 21.5 | *Sphagnum* stems | Poz-172187 | 103.16 ± 0.32 pMC |  |  |
| SM 33.5 | *Sphagnum* stems | Poz-172188 | 104.44 ± 0.33 pMC |  |  |
| SM 43.5 | *Sphagnum* stems | Poz-172189 | 108.12 ± 0.34 pMC |  |  |
