## Supplementary figures and images for "Long-Term Ecological Baselines and Critical Thresholds in Ombrotrophic Peatlands of Europe: Implications for Restoration Strategies"

### Supplementary Figure 1.pdf

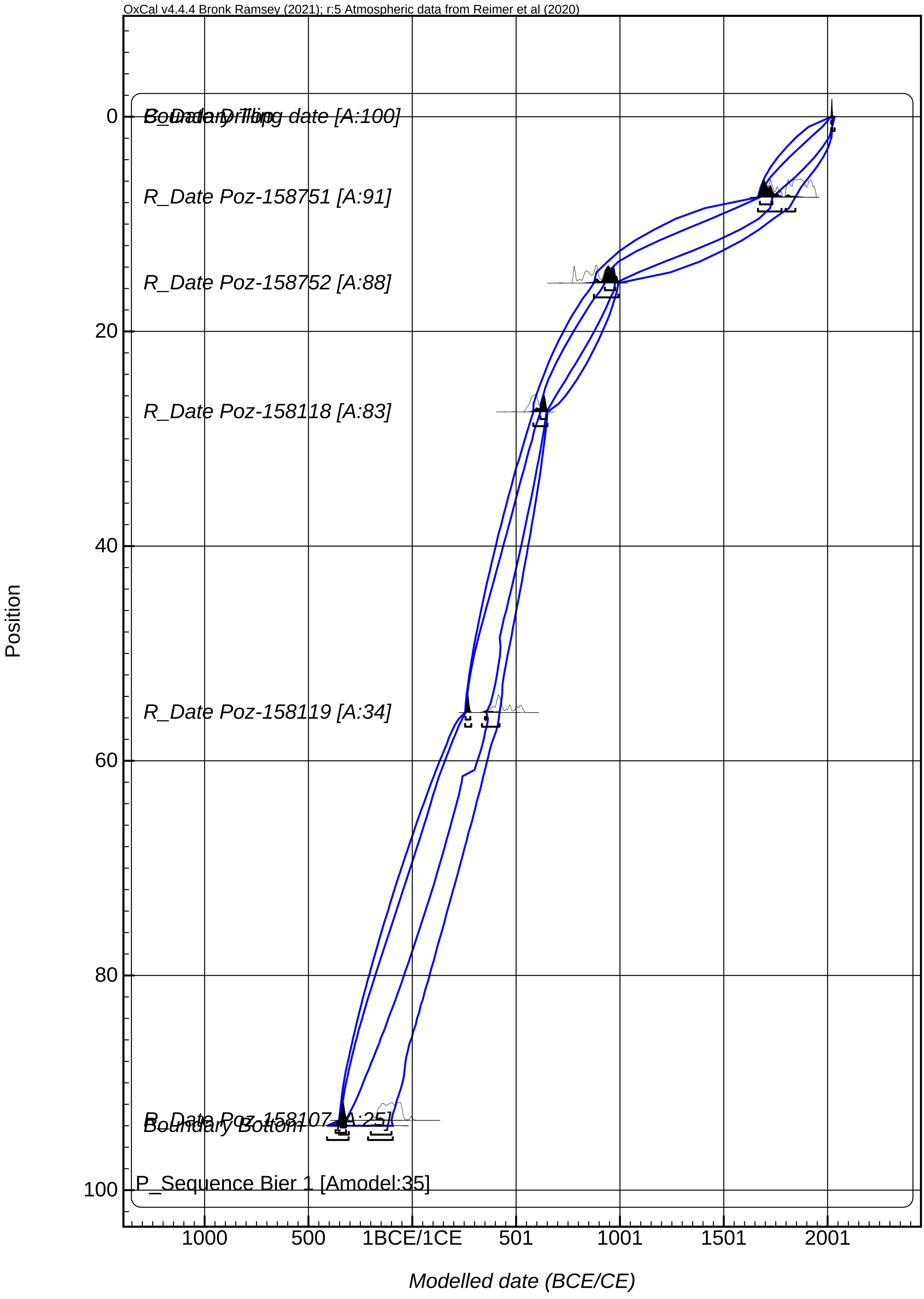

### Supplementary Figure 2.pdf

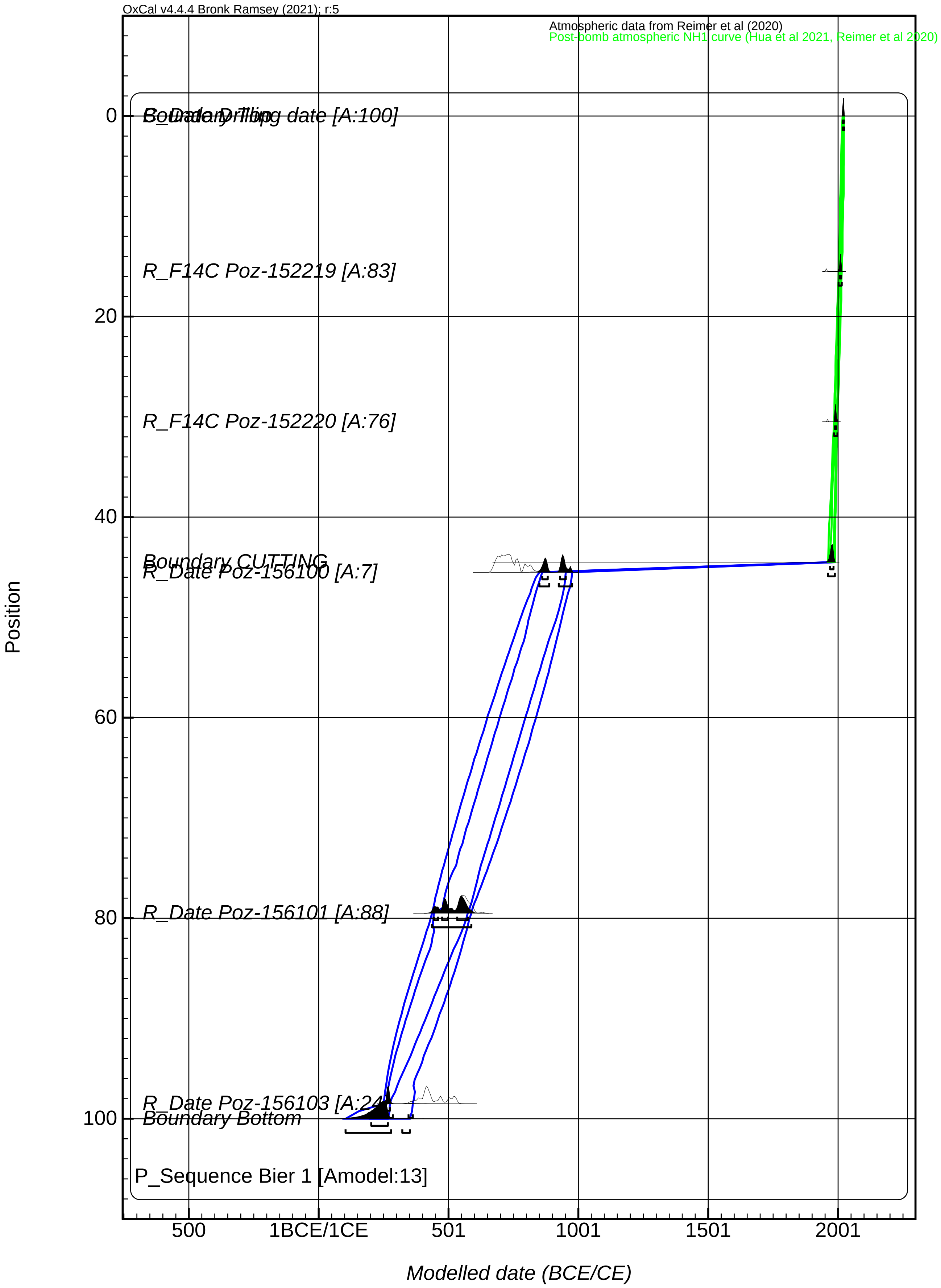

### Supplementary Figure 3.pdf

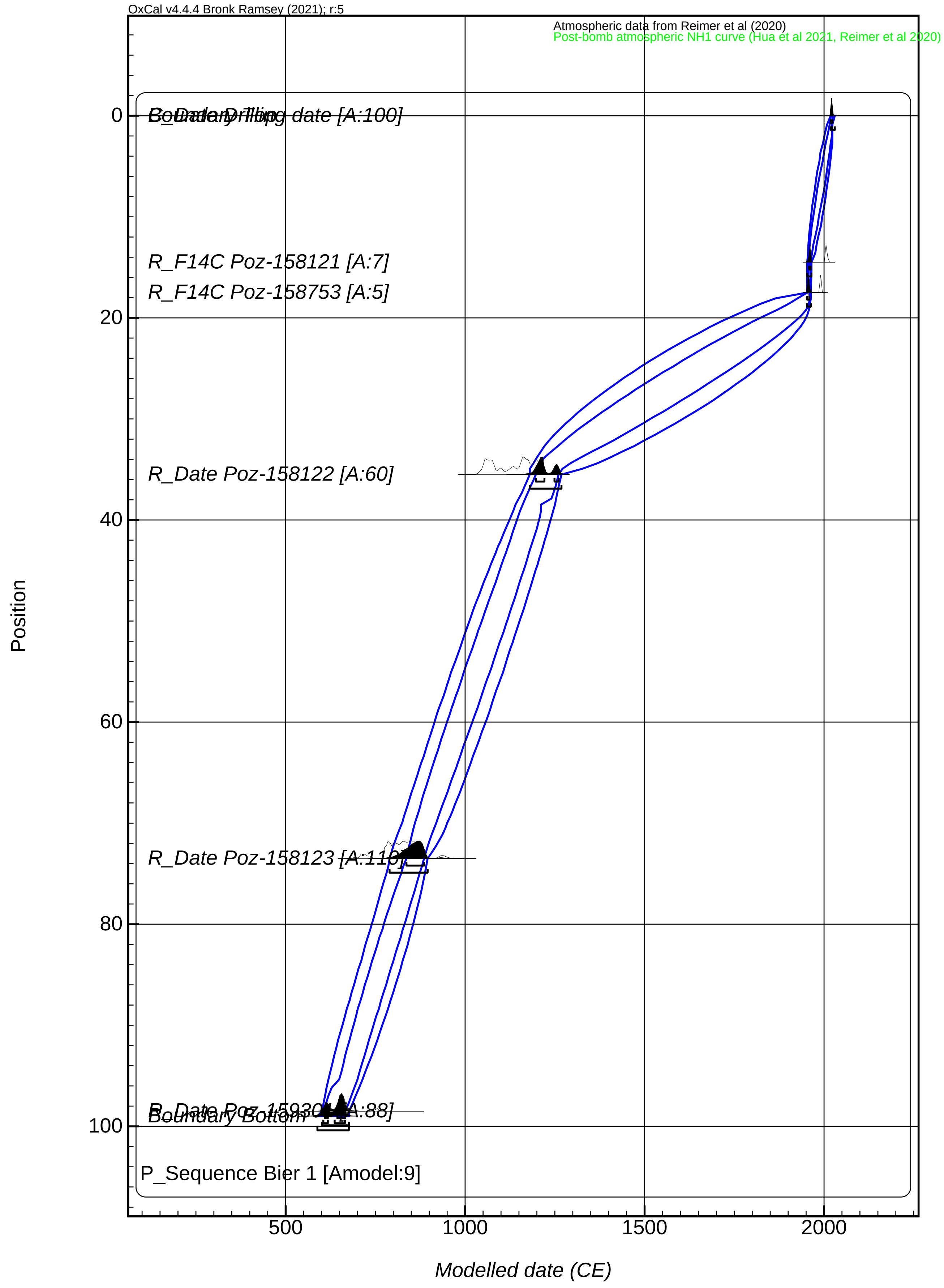

### Supplementary Figure 4.pdf

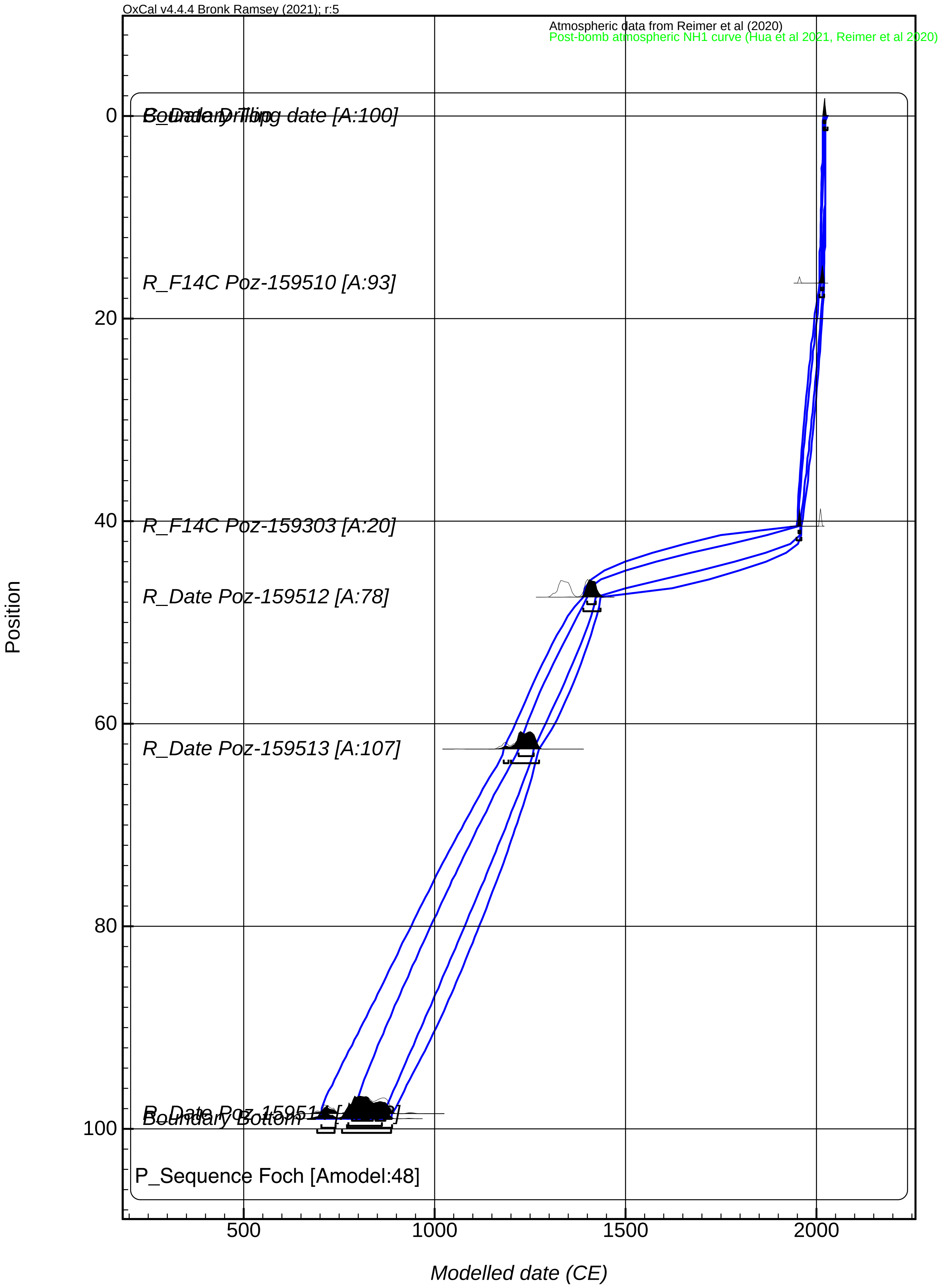

### Supplementary Figure 5.pdf

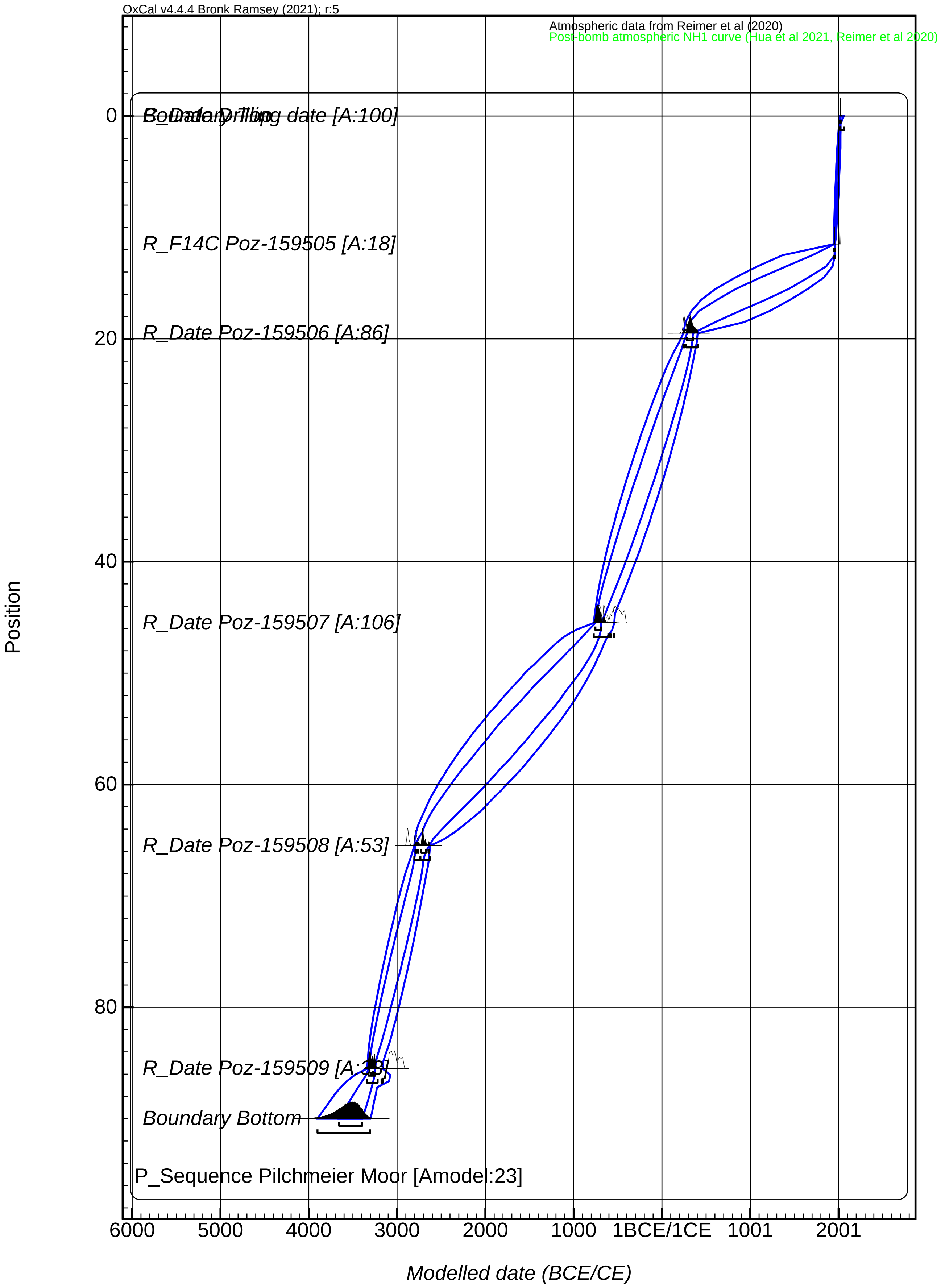

### Supplementary Figure 6.pdf

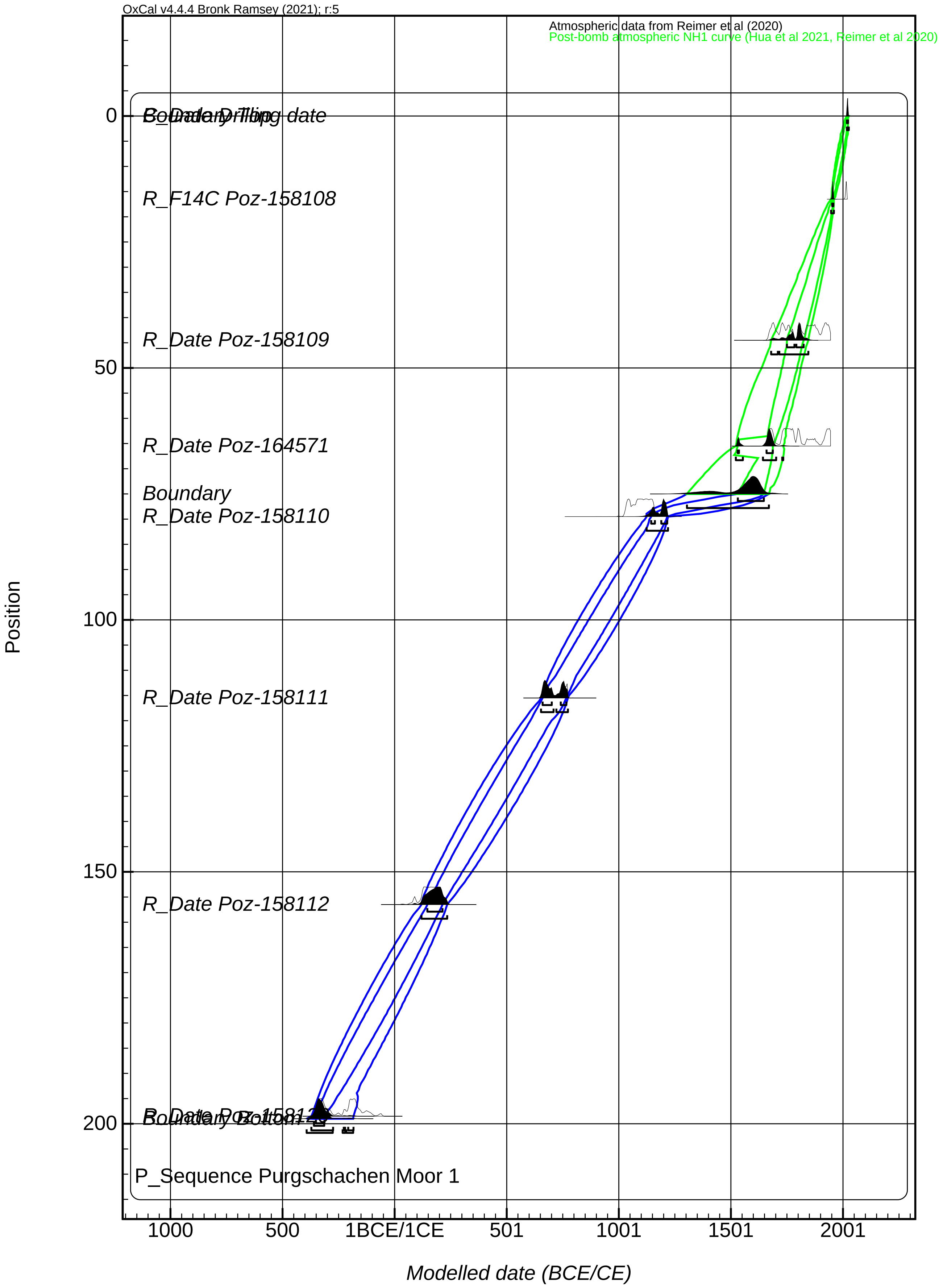

### Supplementary Figure 7.pdf

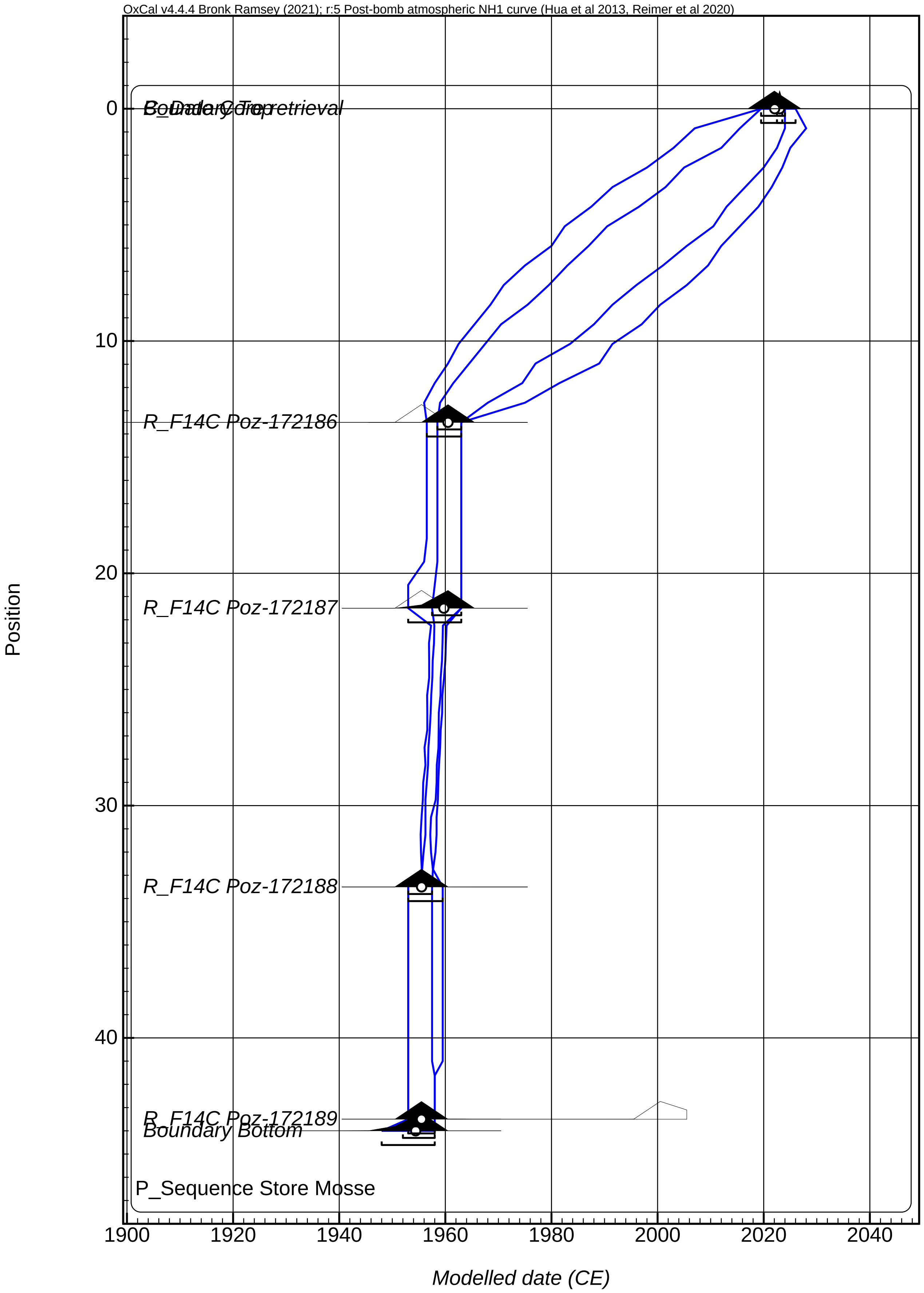

### Supplementary Figure 8.pdf

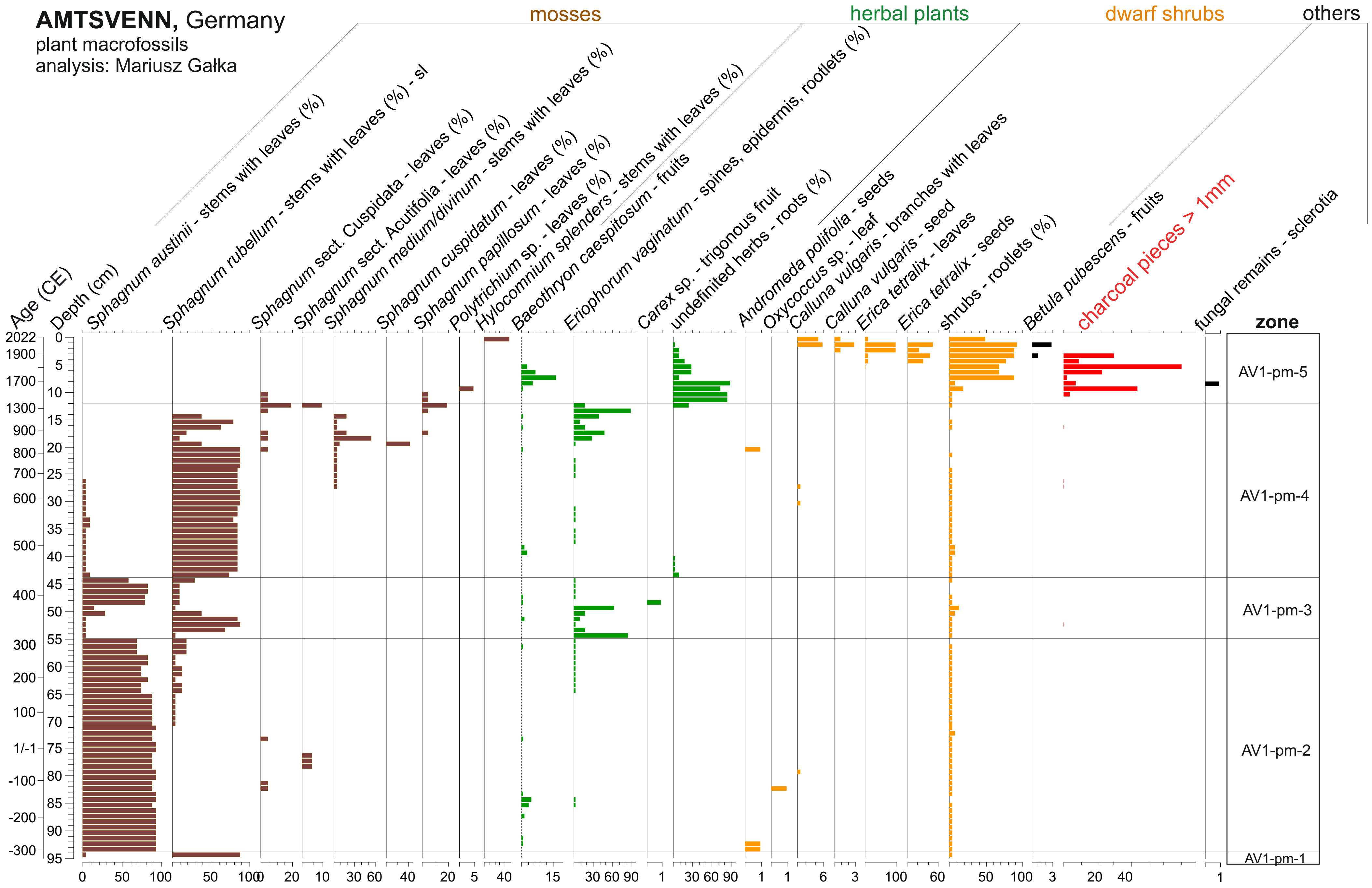

### Supplementary Figure 9.pdf

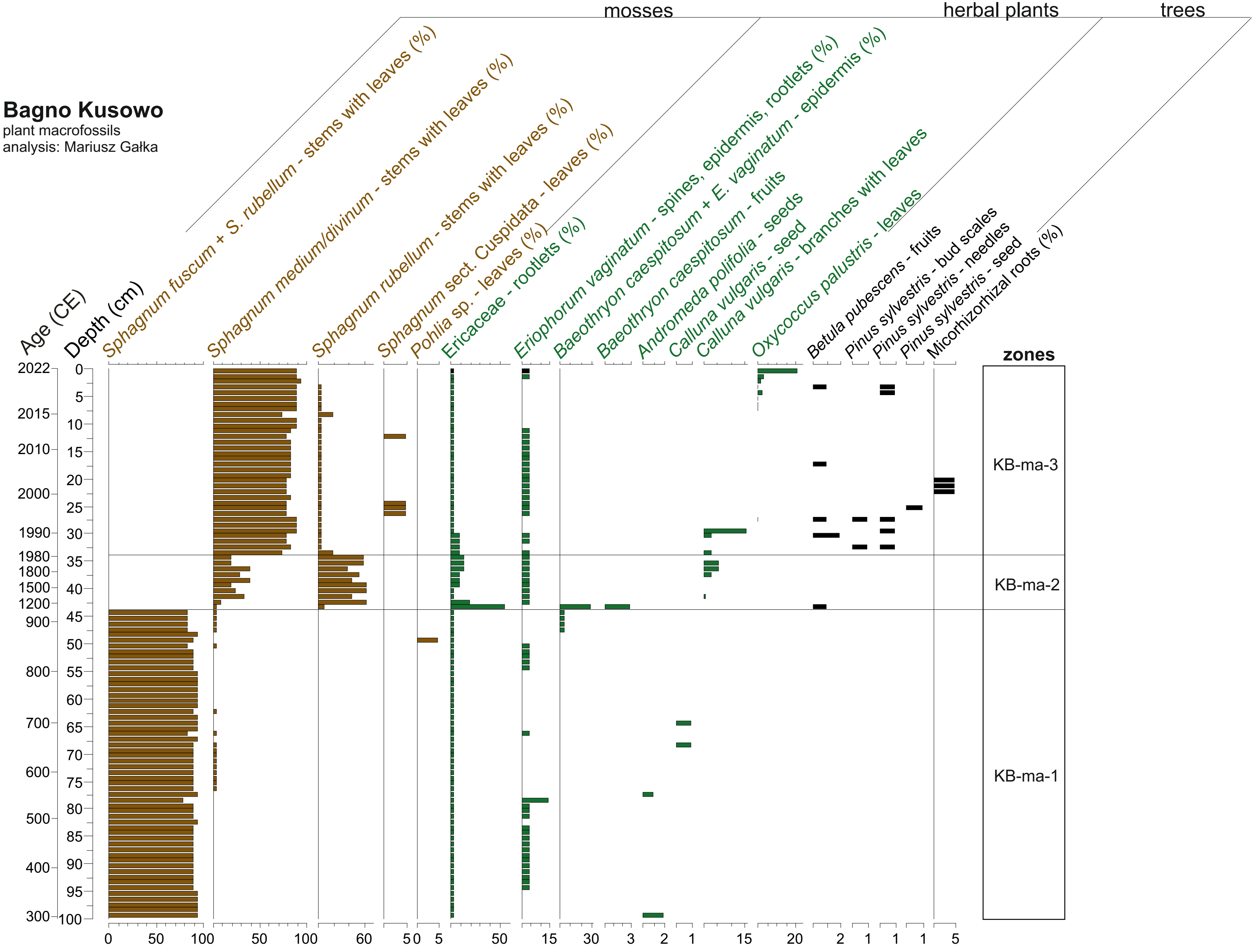

### Supplementary Figure 10.pdf

**Dressersches Moor,** Germany
plant macrofossils
analysis: Mariusz Gałka

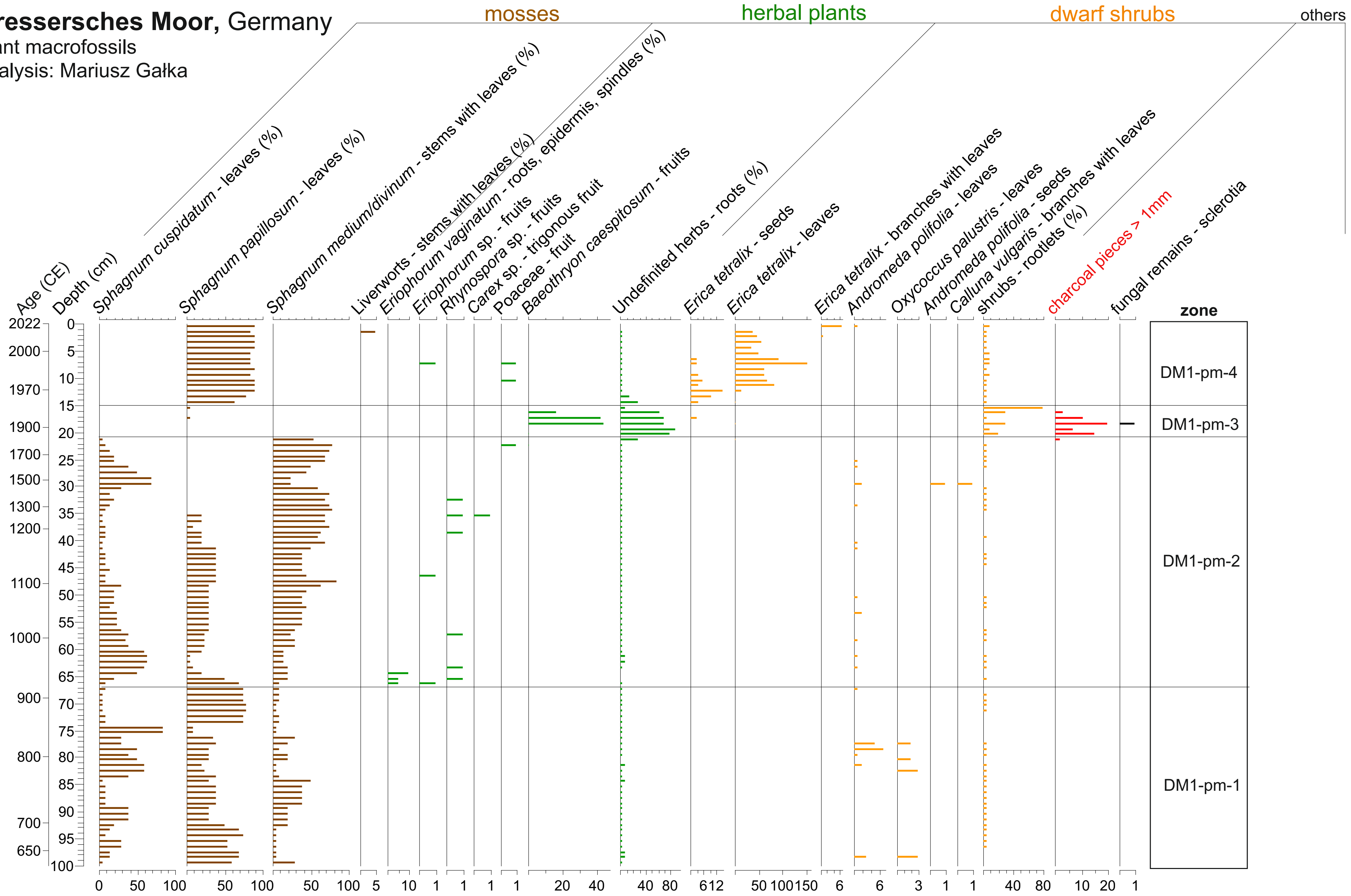

### Supplementary Figure 11.pdf

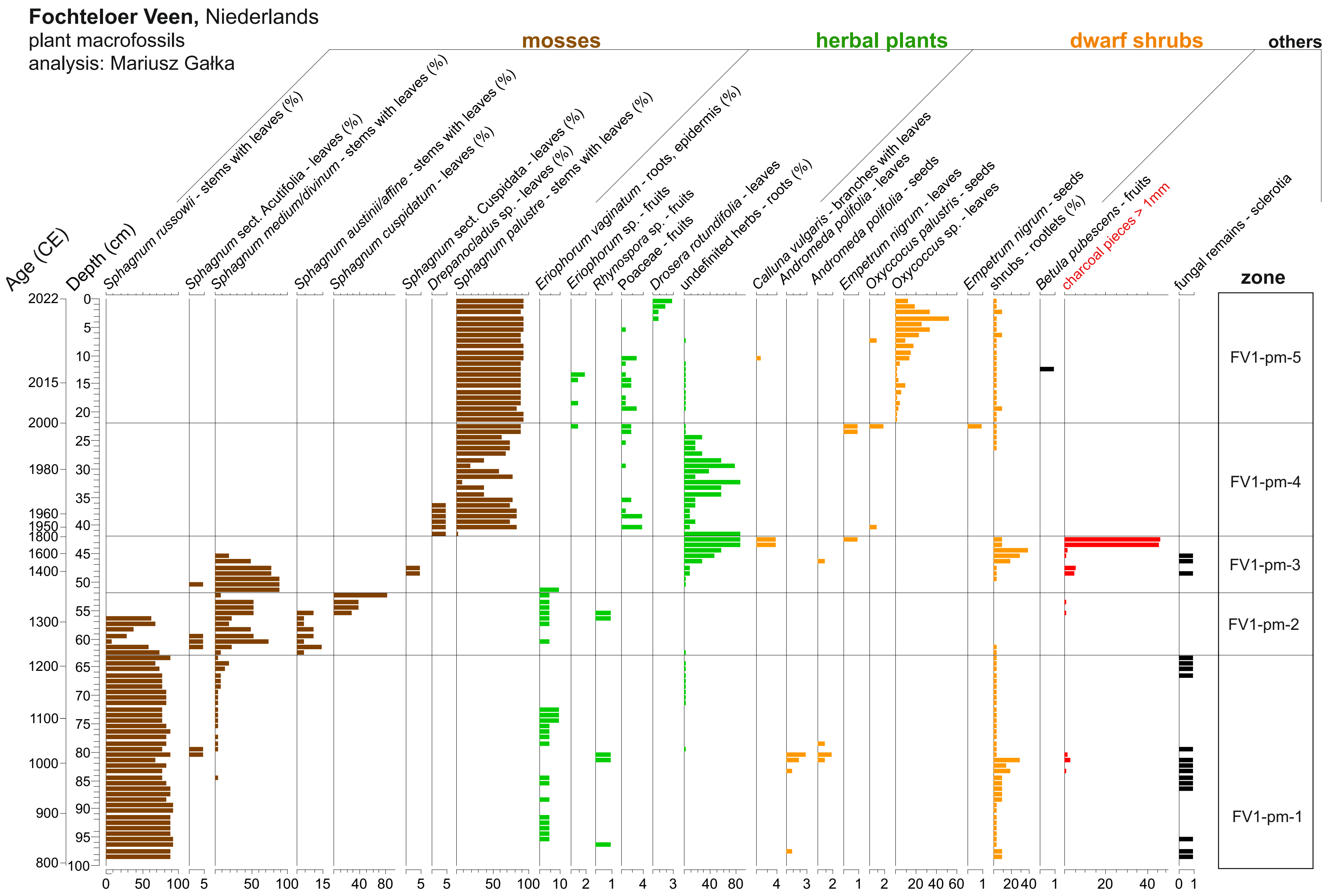

### Supplementary Figure 12.pdf

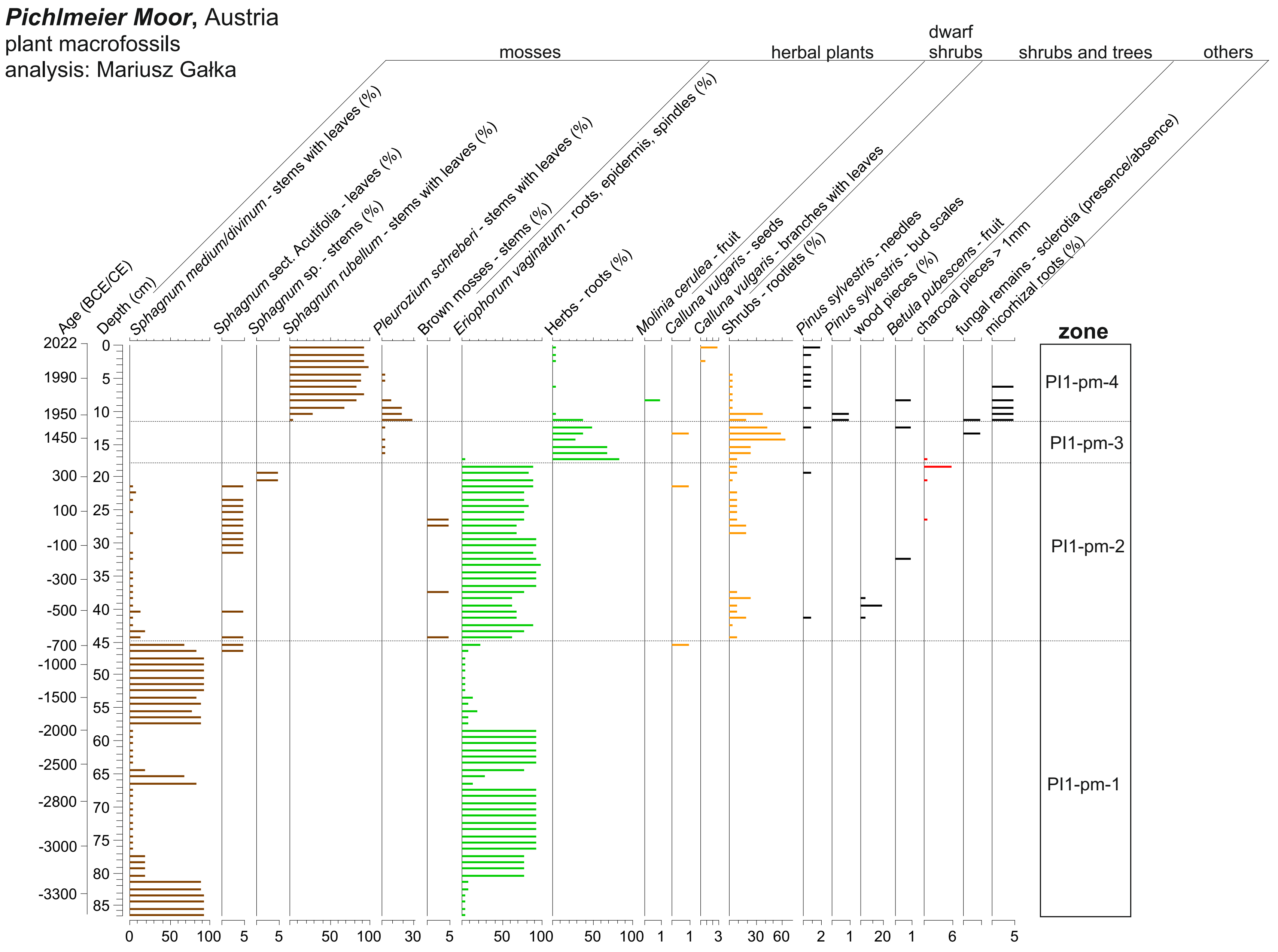

### Supplementary Figure 13.pdf

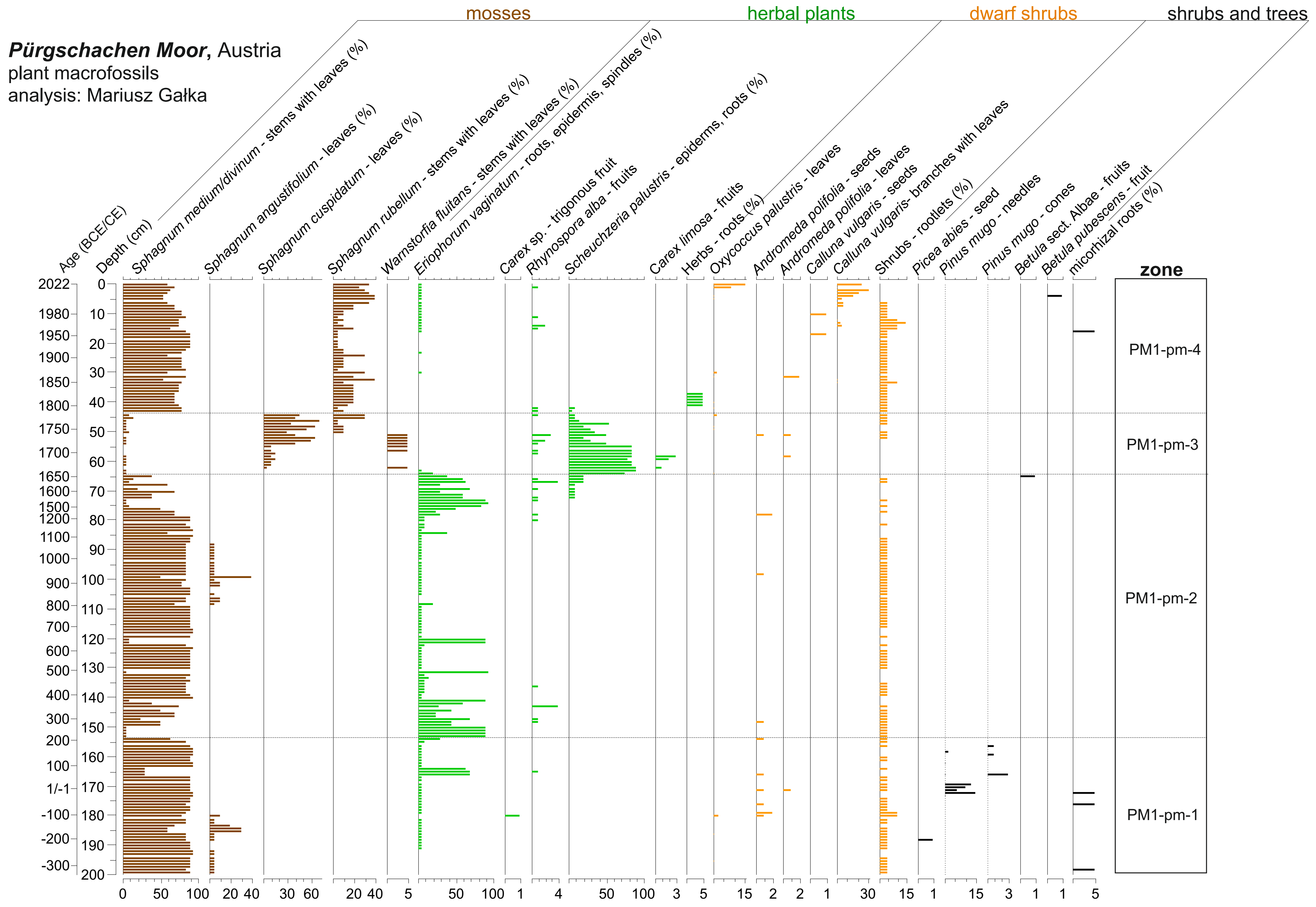

### Supplementary Figure 14.pdf

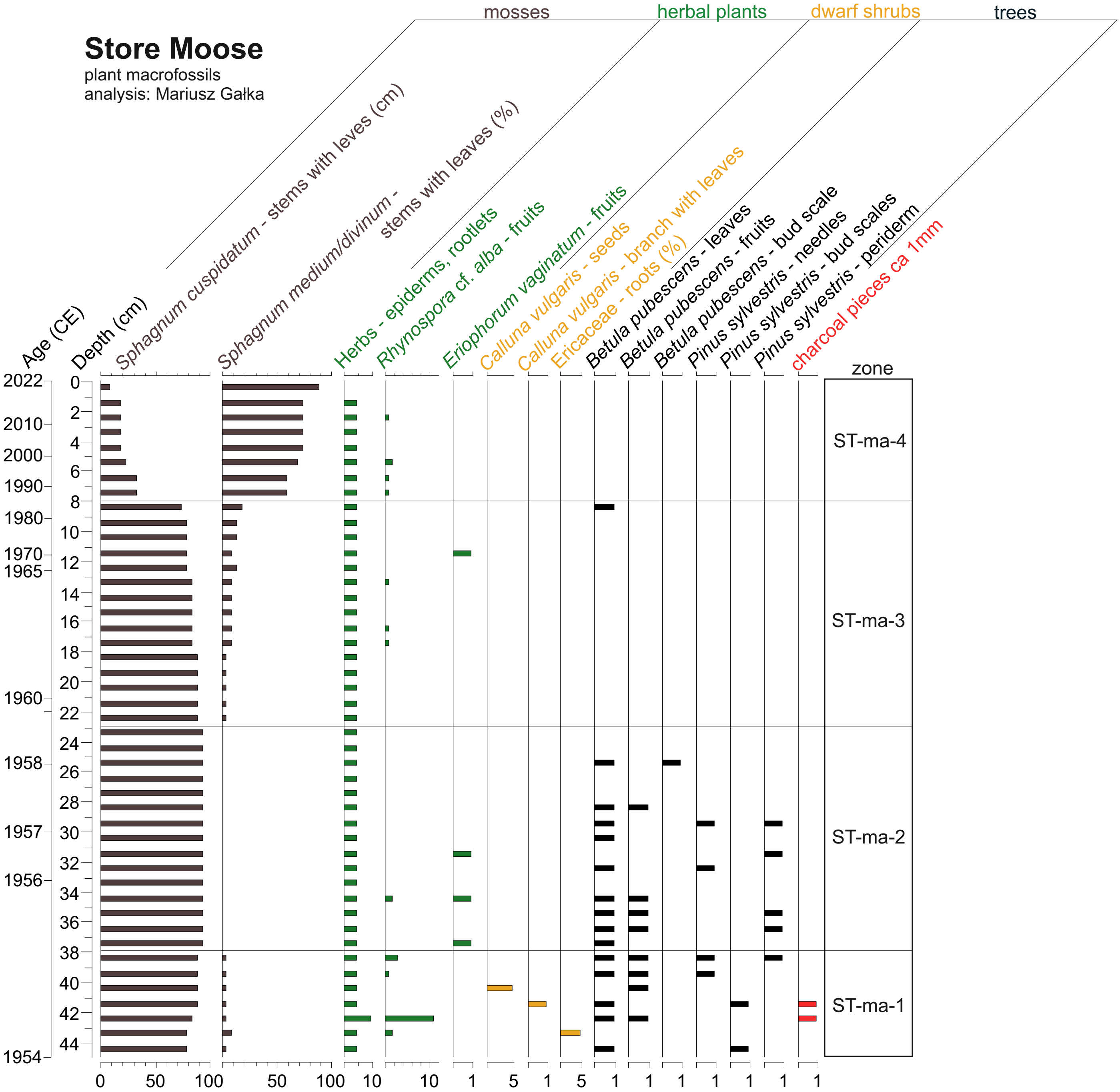

### Supplementary Figure 15.pdf

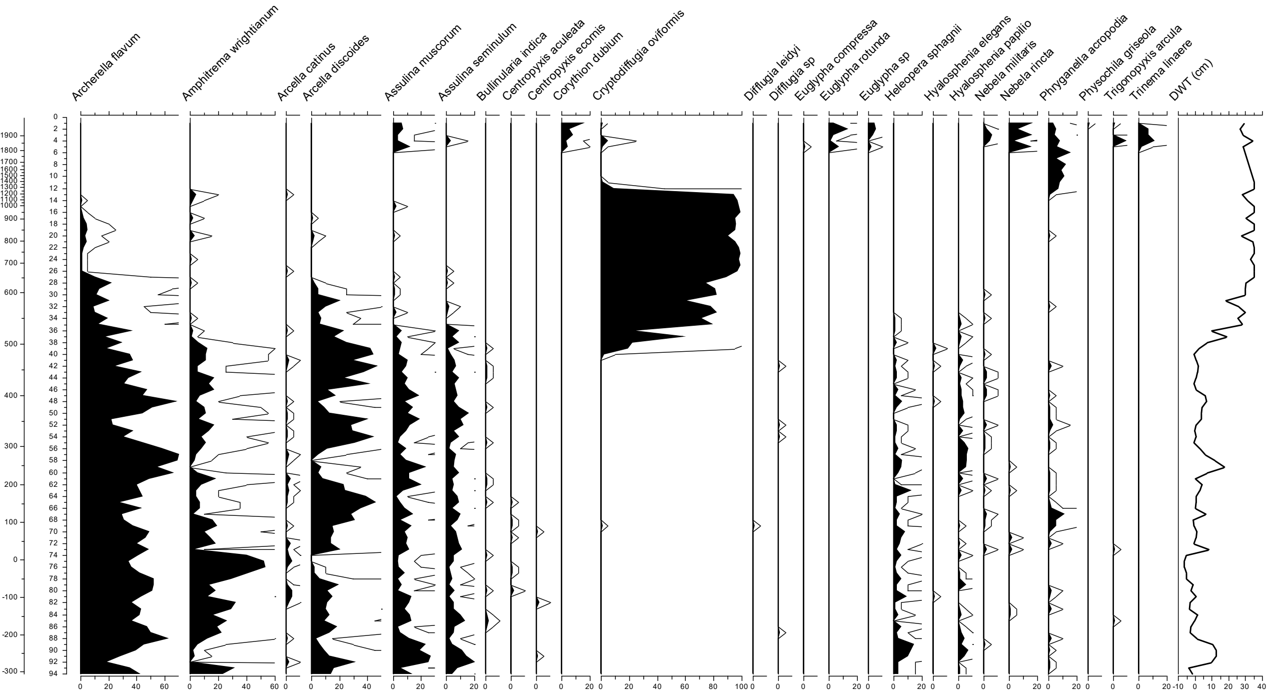

### Supplementary Figure 16.pdf

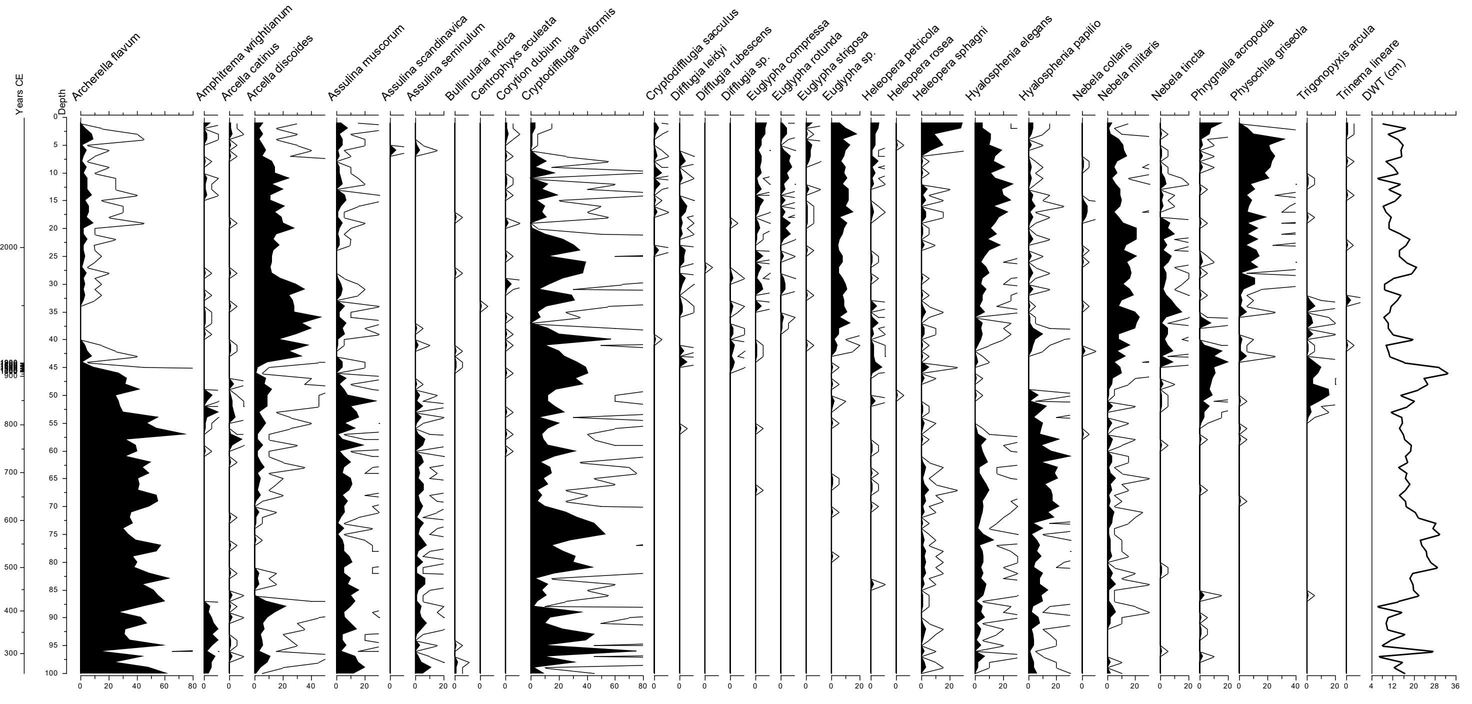

### Supplementary Figure 17.pdf

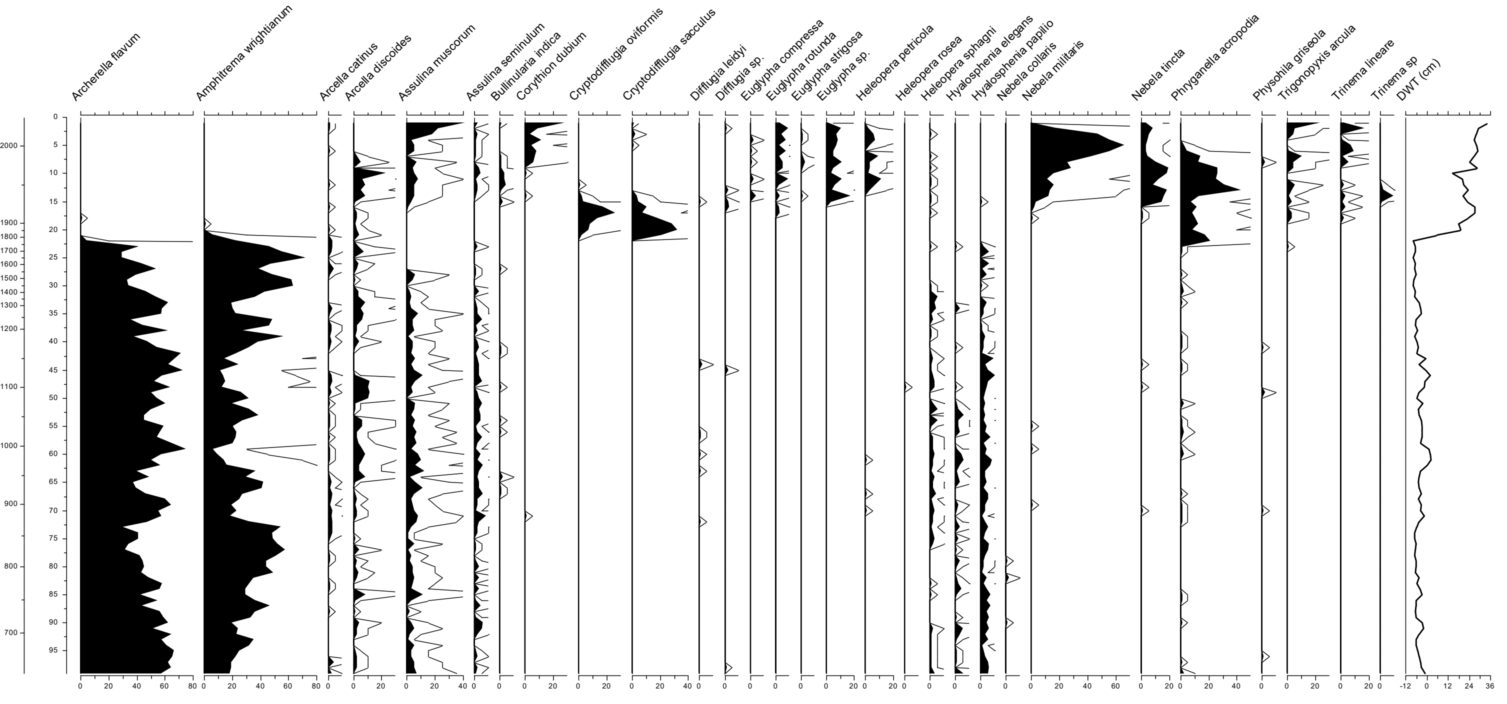

### Supplementary Figure 18.pdf

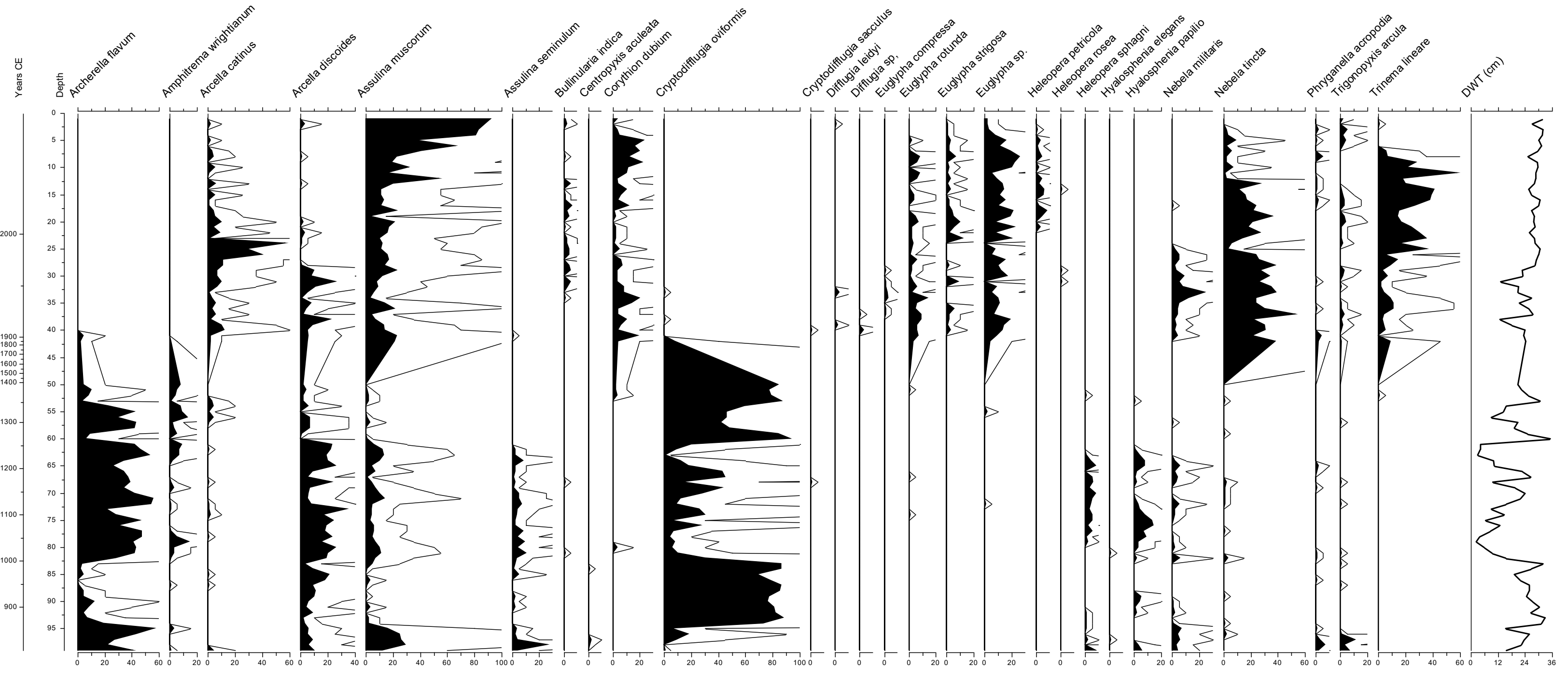

### Supplementary Figure 19.pdf

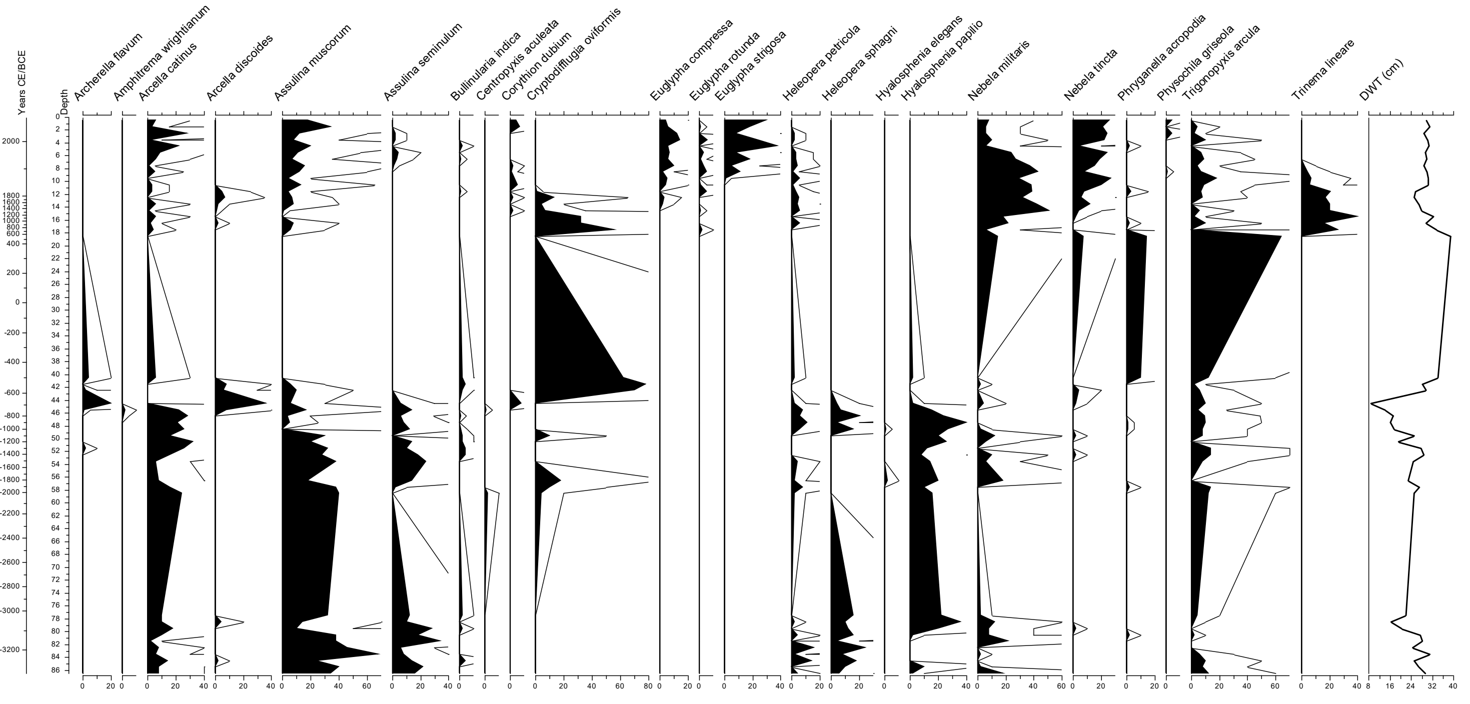

### Supplementary Figure 20.pdf

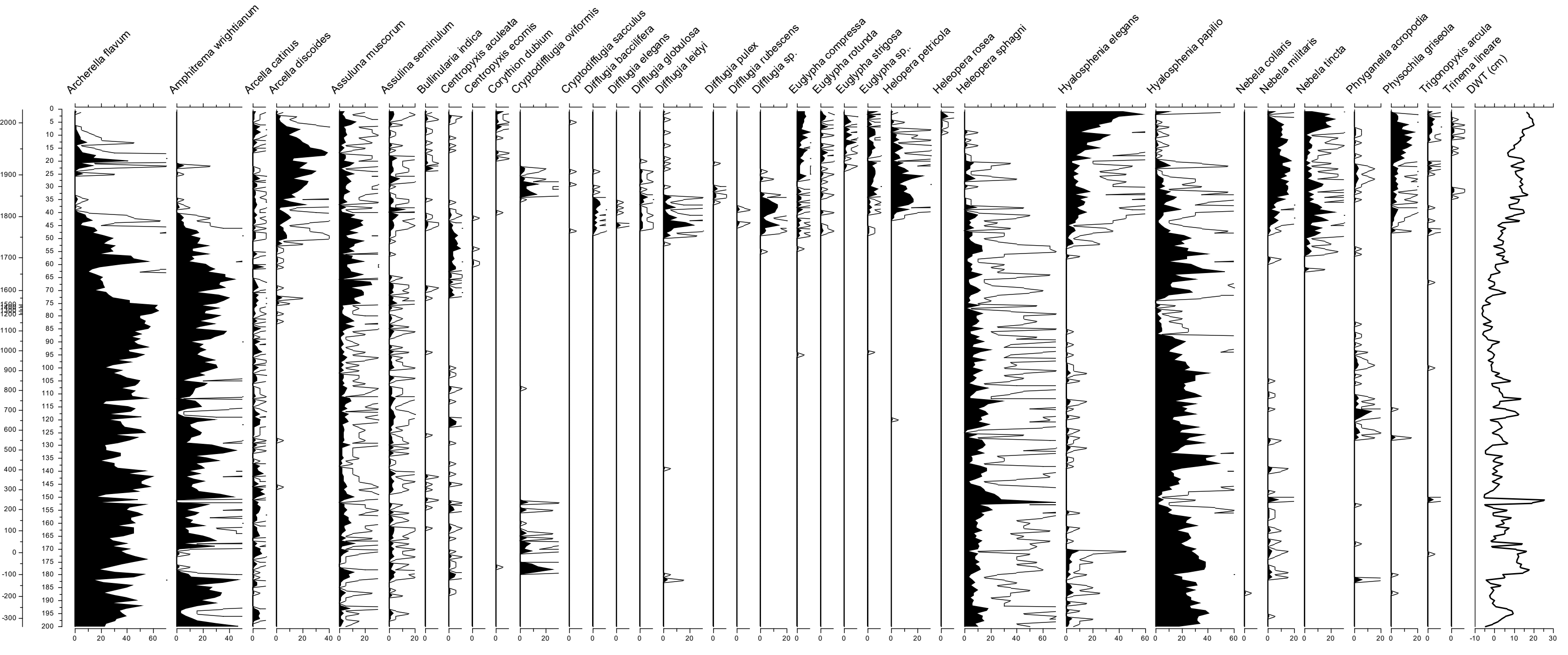

### Supplementary Figure 21.pdf

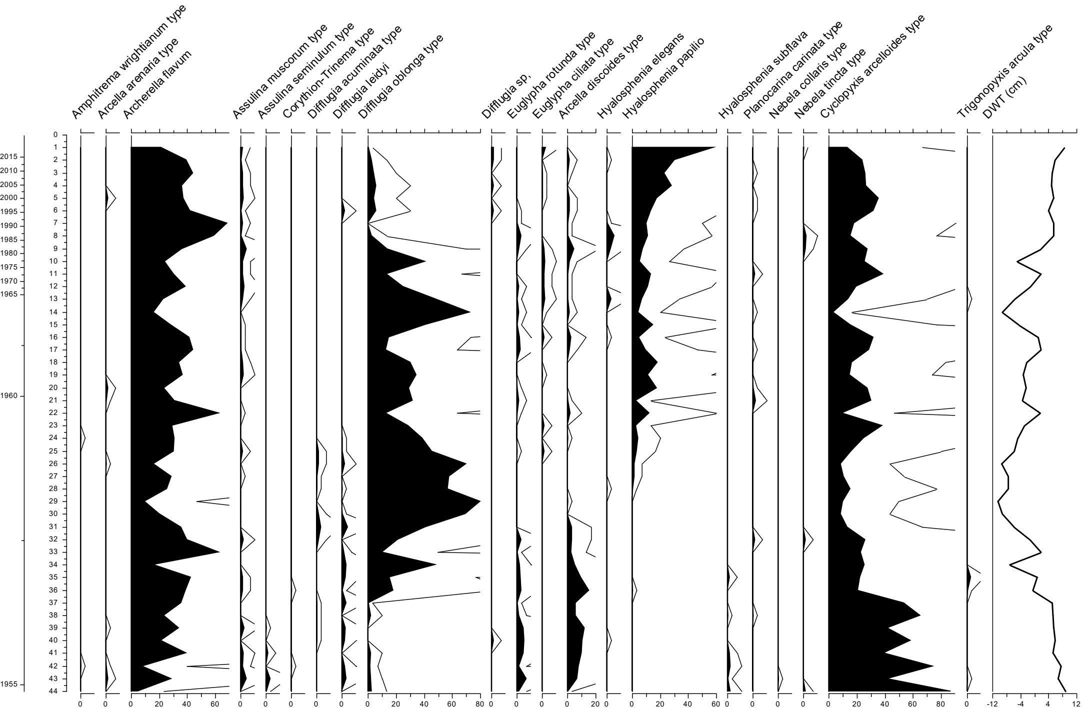
